# Calibration-Aware and Interpretable Graph Learning for Multi-Cohort Diffusion Connectome Brain-Age Modeling

**DOI:** 10.64898/2026.08.19.745783

**Authors:** Alexandra Badea, Ines Poves Acle, Paula Mendez de Inza, Hanwen Lin, Robert J Anderson, Kim G Johnson, Heather E Whitson, Allen W Song, Cristian T Badea, the Alzheimer’s Disease Neuroimaging Initiative, the Health and Aging Brain Study (HABS-HD) Study Team

## Abstract

Brain-age models derived from diffusion MRI–based structural connectomes may provide imaging biomarkers of accelerated brain aging, but their biological interpretation and transportability across heterogeneous populations remain uncertain. We developed a calibration-aware and hierarchically interpretable graph-learning framework and evaluated it across four independent aging and Alzheimer’s disease–related cohorts: ADNI, Duke/UNC ADRC, HABS-HD, and AD-DECODE. The analysis included 1,093 connectome sessions from 789 participants. Cohort-specific graph neural networks were trained using participant-grouped cross-validation across five imaging and multimodal feature configurations. Prediction performance varied more strongly across cohorts than across feature sets, with imaging-only out-of-fold mean absolute error ranging from 4.72 years in ADNI to 9.75 years in AD-DECODE. The imaging-only graph neural network was competitive with ridge, elasticnet, and gradient-boosted regression models trained on matched vectorized connectome features, but was not uniformly superior. Age-bias-corrected brain-age gap was most consistently associated with reduced diffusion-derived microstructural integrity and structural-network organization across cohorts. In longitudinal analyses, corrected brain-age gap showed moderate-to-good within-person preservation in ADNI and HABS-HD, with intraclass correlation coefficients of 0.67 and 0.81, respectively; higher baseline values also predicted subsequent microstructural and network deterioration in ADNI. Multiscale SHAP analysis identified distributed contributions from global graph topology, regional imaging features, edge-derived regional summaries, and individual structural connections involving thalamic, striatal, frontal, parietal, cerebellar, hippocampal, and entorhinal circuitry. External transfer was highly sensitive to cohort shift: across 12 off-diagonal train–test evaluations, median mean absolute error decreased from 17.39 to 8.22 years after target-cohort linear recalibration, whereas median Pearson correlation remained 0.17. Because recalibration used target-cohort chronological age, it was interpreted as a diagnostic sensitivity analysis rather than deployable external validation. Together, these findings support calibration-aware diffusion-connectome brain age as an interpretable imaging biomarker of structural brain aging and prospective microstructural and network vulnerability, while emphasizing the need for cohort-specific calibration before external application.

## I. Introduction

Brain-age modeling estimates an individual’s apparent brain age from neuroimaging data and summarizes the deviation between predicted and chronological age as the brain-age gap (BAG). BAG has emerged as a promising quantitative imaging biomarker because it captures interindividual variation in brain aging beyond chronological age and has been associated with cognitive decline, neurological and psychiatric disease, and neurodegenerative disorders (Cole and Franke 2017) (Gaser, Kalc, and Cole 2024) (Collins et al. 2024) (Van Calster et al. 2019). However, accurate age prediction within a development cohort does not suffice to establish a biologically meaningful or clinically useful biomarker. Translational brain-age models must also address systematic age-related prediction bias, demonstrate biological and longitudinal validity, provide interpretable neuroanatomical information, and maintain acceptable performance when applied beyond the population in which they were developed (de Lange et al. 2022) (Butler et al. 2021) (Van Calster et al. 2019) (Tjoa and Guan 2020; Van Calster et al. 2016) (Sullivan et al. 2015).

Diffusion MRI provides information complementary to conventional structural imaging by characterizing tissue microstructure and structural connectivity, both of which are sensitive to aging and Alzheimer’s disease (Cole 2020) (Beck et al. 2021) (Li et al. 2020). Diffusion-derived connectomes are naturally represented as graphs, with anatomical regions modeled as nodes and interregional connections as edges. Graph neural networks (GNNs) are therefore well suited to connectome-based brain-age modeling because they can integrate regional imaging features, edge attributes, and global network topology within a unified predictive architecture (Kawahara et al. 2017) (Cai, Gao, and Liu 2022) (Gao et al. 2023) (Kazi et al. 2025). Their potential value extends beyond improvements in prediction error: graph-native models can preserve the multiscale organization of the connectome and support interpretation at global, regional, and connection-specific anatomical levels.

Several methodological challenges nevertheless limit the translation of diffusion-connectome brain-age models. Predicted age commonly exhibits regression toward the sample mean, causing raw BAG to remain correlated with chronological age. This residual age dependence can confound downstream associations and produce misleading relationships with age-related clinical or biological variables unless correction is estimated without information leakage (de Lange et al. 2022) (Butler et al. 2021). Predictive performance and corrected BAG should therefore be treated as related but distinct outputs: raw out-of-fold predictions are required for unbiased assessment of age-prediction performance, whereas appropriately corrected BAG is needed for biological and longitudinal interpretation.

Biological validation also remains incomplete in many brain-age studies. Cross-sectional associations with diagnosis, cognition, or a limited set of clinical variables do not establish whether BAG reflects a reproducible participant-specific property of brain aging or whether it identifies vulnerability to subsequent deterioration. A credible imaging biomarker should converge with independent measures of brain integrity, retain sufficient longitudinal stability to distinguish individuals, and remain sensitive to meaningful within-person change. In particular, testing whether baseline BAG predicts subsequent microstructural or network deterioration can help determine whether the measure reflects prospective brain vulnerability rather than residual age-prediction error. The anatomical basis of connectome brain-age predictions is also incompletely understood. Explainable artificial intelligence methods such as SHAP are increasingly used in neuroimaging, but interpretation is often limited to ranked feature importance within a single representation (Lundberg and Lee 2017) (Tjoa and Guan 2020). For graph-based models, a single feature ranking cannot determine whether predictions primarily reflect global network topology, regional tissue properties, the aggregate connectivity of specific regions, or individual structural connections. A hierarchical interpretation strategy spanning these complementary scales is therefore needed to relate graph-model predictions to distributed anatomical and network systems implicated in aging and Alzheimer’s disease, while recognizing that model attributions are descriptive rather than causal.

External transportability presents an additional challenge. Diffusion MRI cohorts can differ substantially in age distribution, demographic composition, vascular and metabolic risk, disease spectrum, scanner hardware, acquisition protocols, preprocessing, and connectome feature distributions. Consequently, models that perform well under within-cohort cross-validation may produce large prediction offsets, compressed or expanded age ranges, or implausible ages when transferred to another cohort. Importantly, external performance reflects at least two distinct properties. Calibration concerns agreement in prediction offset and scale within the target population, whereas discrimination concerns preservation of participant-level age ordering (Van Calster et al. 2016) (Van Calster et al. 2019). A transferred model may retain partial discrimination while being severely miscalibrated, and post-hoc recalibration may improve absolute agreement without recovering lost participant-level information. These dimensions should therefore be examined separately rather than summarized by a single aggregate performance metric.

Prediction accuracy, age-bias correction, biological validation, longitudinal robustness, interpretability, calibration, and external transportability are commonly evaluated in isolation. This fragmented approach makes it difficult to determine whether a brain-age model constitutes a credible imaging biomarker or merely performs adequately as an age regressor within its development cohort. A unified evaluation framework is needed to examine these complementary dimensions together.

In this study, we developed a calibration-aware and hierarchically interpretable graph-learning framework for diffusion-connectome brain-age modeling across four heterogeneous aging and Alzheimer’s disease–related cohorts: ADNI, Duke/UNC ADRC, HABS-HD, and AD-DECODE. The analysis included 1,093 diffusion MRI/connectome sessions from 789 participants with distinct demographic, vascular, metabolic, clinical, and molecular profiles. Structural connectomes were represented as graphs, and cohort-specific GNNs were trained using participant-grouped cross-validation to prevent leakage across repeated sessions. Five feature configurations were evaluated, spanning imaging-only and multimodal inputs. Raw out-of-fold predicted age was retained for primary predictive-performance assessment, whereas age-bias-corrected brain-age gap (cBAG) was used for downstream biological and longitudinal validation.

The principal contributions of this work are fivefold. First, we provide a leakage-safe, multi-cohort evaluation of diffusion-connectome brain-age prediction using graph-native representations and benchmark the imaging-only GNN against conventional regression models trained on matched vectorized connectome features. Second, we evaluate cBAG as an imaging biomarker through cross-sectional biological associations, longitudinal preservation of individual brain-aging position, and prediction of subsequent microstructural and network deterioration. Third, we introduce a multiscale SHAP framework integrating global graph measures, node-level features, edge-derived regional summaries, and individual structural connections. Fourth, we systematically characterize external train–test transfer across cohorts and use post-hoc recalibration sensitivity analyses to distinguish calibration mismatch from loss of participant-level discrimination. Finally, we integrate prediction, biological validity, longitudinal robustness, interpretability, calibration, and external transportability within a unified evaluation framework. Together, these analyses examine whether diffusion-connectome brain age can serve as an interpretable imaging biomarker of structural brain aging and prospective network vulnerability across heterogeneous populations.

## II. Methods

### A. Study Cohorts and Harmonized Inputs

This study used diffusion MRI/connectome data from four independent aging and Alzheimer’s disease–related cohorts: ADNI, Duke/UNC ADRC, HABS-HD, and AD-DECODE. Data from the Alzheimer’s Disease Neuroimaging Initiative (ADNI) were obtained from the ADNI database. ADNI is a longitudinal public–private study established to evaluate imaging, biomarker, clinical, and neuropsychological measures of Alzheimer’s disease progression and to support biomarker validation for clinical trials. Data used in this study were downloaded on February 5, 2025, and checked for updates before manuscript submission. Participants were included if they had usable diffusion MRI/connectome data and the required demographic and analysis-specific information were available. Participant identifiers were retained throughout cross-validation to prevent data from the same individual from entering both training and held-out folds.

Data from the Health and Aging Brain Study–Health Disparities (HABS-HD), a cohort enriched for metabolic and cardiometabolic risk, were accessed on February 5, 2025 (Petersen et al. 2025). A deidentified dataset from the Duke/UNC Alzheimer’s Disease Research Center (ADRC) was obtained on June 14, 2025. The Duke/UNC ADRC sample included adults spanning approximately 28–80 years of age and was enriched for cognitively unimpaired individuals with a family history of dementia. After metadata harmonization, the final dataset comprised 1,093 diffusion MRI/connectome sessions from 789 unique participants: 316 sessions from 180 ADNI participants, 204 sessions from 204 Duke/UNC ADRC participants, 487 sessions from 319 HABS-HD participants, and 86 sessions from 86 AD-DECODE participants. ADNI and HABS-HD contributed repeated-session data, whereas Duke/UNC ADRC and AD-DECODE were analyzed as cross-sectional cohorts for primary model-performance evaluation. **Table 1** summarizes demographic, clinical, genetic, and cognitive characteristics across cohorts.

**Table 1.** Cohort characteristics and graph-aligned data availability. Sample size is reported as both connectome sessions and participants because ADNI and HABS-HD contributed longitudinal data, whereas Duke/UNC ADRC and AD-DECODE were analyzed cross-sectionally. APOE ε4 carriage was defined by APOE3/4 or APOE4/4 status and excluded APOE2/4. Participants were included only when the required connectome data, harmonized identifiers, and analysis-specific covariates or outcomes were available. Abbreviations: AD, Alzheimer’s disease; ADNI, Alzheimer’s Disease Neuroimaging Initiative; Duke/UNC ADRC, Duke/University of North Carolina Alzheimer’s Disease Research Center; APOE, apolipoprotein E; DTI, diffusion tensor imaging; FA, fractional anisotropy; HABS-HD, Health and Aging Brain Study–Health Disparities; MCI, mild cognitive impairment; SD, standard deviation.

| Characteristic | ADNI | Duke/UNC ADRC | HABS-HD | AD-DECODE |
| --- | --- | --- | --- | --- |
| **Cohort information** |  |  |  |  |
| Data accessed | Longitudinal | Cross-sectional | Longitudinal | Cross-sectional |
| Modalities/features included | DTI connectome; FA; volume; graph metrics; APOE; vascular measures | DTI connectome; FA; volume; graph metrics; APOE; vascular measures | DTI connectome; FA; volume; graph metrics; APOE; vascular measures | DTI connectome; FA; volume; graph metrics; APOE; vascular measures |
| **Sample size** |  |  |  |  |
| Connectome/DWI sessions, n | 316 | 204 | 487 | 86 |
| Participants, n | 180 | 204 | 319 | 86 |
| **Demographic characteristics** |  |  |  |  |
| Age, years, mean ± SD | 75.97 ± 5.43 | 57.96 ± 11.85 | 66.60 ± 8.55 | 52.18 ± 16.12 |
| Age, years, median [IQR] | 75.45 [71.75, 78.93] | 58.00 [50.00, 67.00] | 66.67 [60.35, 72.65] | 51.00 [40.00, 66.38] |
| Age, years, range | 67.3–93.4 | 28.0–80.0 | 50.0–90.2 | 20.2–83.0 |
| Female, n (%) | 90 (50.0%) | 140 (68.6%) | 216 (67.7%) | 47 (54.7%) |
| Male, n (%) | 89 (49.4%) | 64 (31.4%) | 103 (32.3%) | 39 (45.3%) |
| **APOE genotype** |  |  |  |  |
| APOE ε4 carriers (%) | 62 (35.2%) | 88 (48.9%) | 86 (28.5%) | 39 (45.3%) |
| APOE22 n (%) | 0 (0.0%) | 0 (0.0%) | 2 (0.6%) | 0 (0.0%) |
| APOE23 n (%) | 18 (10.0%) | 18 (8.8%) | 29 (9.1%) | 8 (9.3%) |
| APOE24 n (%) | 4 (2.2%) | 5 (2.5%) | 9 (2.8%) | 0 (0.0%) |
| APOE33 n (%) | 96 (53.3%) | 74 (36.3%) | 185 (58.0%) | 39 (45.3%) |
| APOE34 n (%) | 55 (30.6%) | 71 (34.8%) | 76 (23.8%) | 31 (36.0%) |
| APOE44 n (%) | 7 (3.9%) | 17 (8.3%) | 10 (3.1%) | 8 (9.3%) |
| **Clinical/diagnostic group** |  |  |  |  |
| Cognitively normal/normal, n (%) | 134 (74.4%) | 145 (71.1%) | 254 (79.6%) | 79 (91.9%) |
| MCI, n (%) | 41 (22.8%) | 16 (7.8%) | 56 (17.6%) | 5 (5.8%) |
| AD/dementia, n (%) | 5 (2.8%) | 9 (4.4%) | 8 (2.5%) | 2 (2.3%) |
| **Cognitive composites** |  |  |  |  |
| Global cognition composite, n (%) | 180 (100.0%) | 198 (97.1%) | 318 (99.7%) | 86 (100.0%) |
| Global cognition composite, mean ± SD | 0.05 ± 0.74 | -0.00 ± 1.00 | 0.02 ± 0.59 | -0.00 ± 1.01 |

The cohorts differed in age, sex distribution, diagnostic composition, and APOE genotype availability. ADNI was the oldest cohort (75.97 ± 5.43 years), followed by HABS-HD (66.60 ± 8.55 years), Duke/UNC ADRC (57.96 ± 11.85 years), and AD-DECODE (52.18 ± 16.12 years). Females represented 50.0% of ADNI, 68.6% of Duke/UNC ADRC, 67.7% of HABS-HD, and 54.7% of AD-DECODE. Cognitively normal participants constituted the majority of each cohort, whereas MCI and AD/dementia cases were present at lower frequencies. Strict APOE ε4 carriage, defined as APOE3/4 or APOE4/4 and excluding APOE2/4, ranged from 28.5% in HABS-HD to 48.9% in Duke/UNC ADRC among participants with classifiable genotype data. Global cognitive composite scores were available for nearly all participants across cohorts. For each cohort, harmonized metadata were used to define participant identifiers, session identifiers, chronological age, sex, diagnostic group, APOE genotype, cognitive variables, vascular measures, and available biomarker or transcriptomic variables. Diagnostic labels were harmonized into cognitively normal, mild cognitive impairment, AD/dementia, and other or unknown categories when available. Primary model training and out-of-fold evaluation were performed within the cognitively normal training sample. Full-cohort prediction outputs were generated separately for downstream descriptive and validation analyses.

MRI data were acquired according to cohort-specific protocols. ADNI comprised heterogeneous 3-T multisite acquisitions across Siemens, GE, and Philips platforms, including predominantly single-shell diffusion imaging at b = 1000 s/mm² and a subset of multishell acquisitions. Duke/UNC ADRC used a uniform 3-T GE single-shell diffusion protocol with 25 diffusion-weighted directions at b = 800 s/mm² and two b = 0 volumes. HABS-HD used a standardized 3-T Siemens protocol with 192 diffusion-weighted volumes at b = 1000 s/mm² and 12 b = 0 volumes. AD-DECODE used a 3-T GE four-shot MUSE diffusion protocol with 21 directions at b = 1000 s/mm² and three b = 0 volumes. Anatomical imaging was acquired in all cohorts using cohort-specific 3D T1-weighted protocols. Image acquisition details are provided in **Supplementary Table S1.** The analysis workflow linked cohort-specific diffusion/connectome graph inputs to harmonized metadata, feature-set construction, graph neural network training, raw out-of-fold age prediction, age-bias correction, and validation (**Fig. 1**).

**Figure 1.**
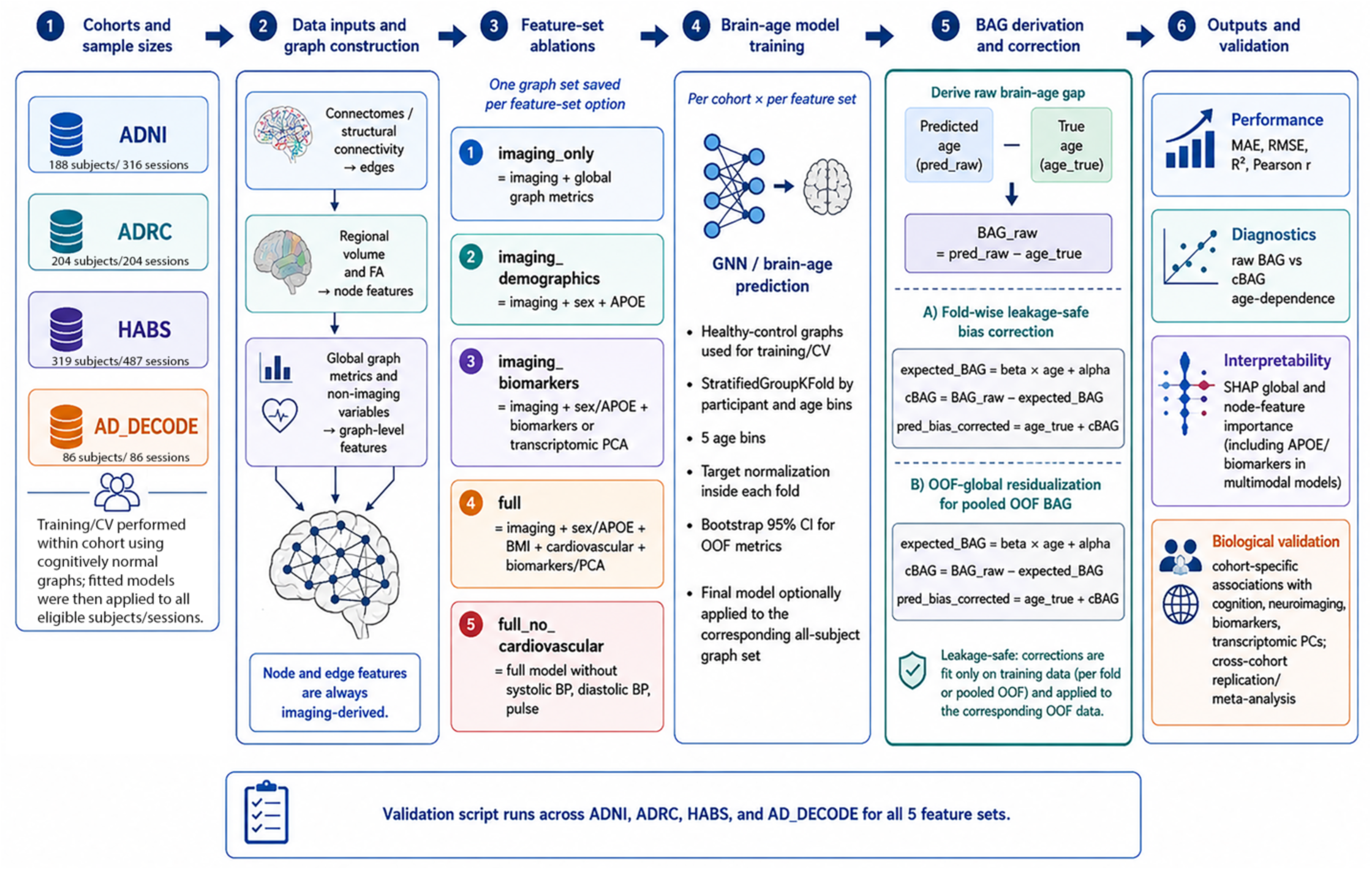
The brain-age modeling pipeline. Diffusion MRI/connectome graphs from ADNI, Duke/UNC ADRC, HABS-HD, and AD-DECODE were aligned to harmonized cohort metadata. Brain regions were represented as nodes and connections as edges. Cohort-specific imaging, graph-theoretic, demographic, genetic, vascular, biomarker, and transcriptomic variables were organized into predefined feature sets according to data availability. Brain-age models were trained separately within each cohort and feature set using participant-grouped cross-validation in cognitively normal samples. Raw out-of-fold predicted age was used to quantify prediction performance, whereas age-bias-corrected brain-age gap (cBAG) was used for downstream validation. Outputs were evaluated using prediction-performance metrics, age-bias diagnostics, SHAP interpretability, biological association analyses, longitudinal, rank-stability, and cross-cohort transferability analyses.

### B. Diffusion Connectome Graph Construction

Diffusion MRI data were processed using an MRtrix3-based tractography pipeline. Diffusion gradient orientations were checked using dwigradcheck, with the required gradient-axis transformations applied before subsequent processing. Anatomical and diffusion images were converted to MRtrix image format, and diffusion tensor metrics, including fractional anisotropy (FA), mean diffusivity, axial diffusivity, and radial diffusivity, were derived using dwi2tensor and tensor2metric. Subject-specific anatomical label images were generated using SAMBA, with cerebrospinal-fluid regions excluded for tractography masking. Multi-tissue response functions for white matter, gray matter, and cerebrospinal fluid were estimated using the Dhollander algorithm, followed by multi-tissue constrained spherical deconvolution and intensity normalization of the resulting fiber-orientation distributions. Whole-brain probabilistic tractography was performed from the gray–white matter interface, initially generating 10 million streamlines per session using an FA cutoff of 0.1 and a maximum streamline length of 1000 mm. A subset of 2 million streamlines was retained for connectome construction. Structural connectomes were defined using an anatomical parcellation comprising 84 regions. Pairwise structural connectivity was quantified using the number of reconstructed streamlines connecting each pair of regions. Symmetric streamline-count connectivity matrices with zero-valued diagonals were generated using tck2connectome. Although additional connectome variants were generated during preprocessing, the plain streamline-count connectome was used for the brain-age graph models. For graph neural network analyses, anatomical regions were represented as graph nodes and nonzero pairwise structural connections as graph edges. Edge attributes encoded streamline-count connectivity between regions. Node-level imaging features included regional measures such as FA and regional volume, while graph-level imaging features included clustering coefficient, characteristic path length, local efficiency, and global efficiency. Session-level graphs were linked to harmonized participant metadata using cohort-specific graph identifiers, and graph-coverage quality-control checks were performed before model training and downstream analyses

### C. Feature-Set Configurations

Five feature-set configurations were evaluated. The imaging-only model included diffusion/connectome graph inputs and imaging-derived graph-level features. The imaging plus demographics model additionally included sex and APOE genotype. The imaging plus biomarkers model additionally included available cohort-specific biomarker variables or transcriptomic principal components. The full model included imaging-derived graph features, sex, APOE genotype, body mass index, cardiovascular variables, and available biomarker or transcriptomic features. The full model without cardiovascular variables used the same multimodal feature set as the full model but excluded systolic blood pressure, diastolic blood pressure, and pulse. This feature-ablation design was used to assess whether adding non-imaging covariate blocks improved brain-age prediction beyond imaging-derived graph information. Because biomarker and transcriptomic variables differed across cohorts, feature matrices were constructed separately within cohort using the harmonized variables available for that cohort. For conventional baseline comparisons, the imaging-only graph inputs were converted to a vector comprising vectorized connectome edge attributes, global graph metrics, and summary statistics of node-level features. For each node-feature channel, the mean, standard deviation, minimum, and maximum across brain regions were appended to the feature vector, yielding 3,502 features per participant/session. Three conventional regression models were evaluated using this representation: Ridge regression (RidgeCV), ElasticNet regression (ElasticNetCV), and extreme gradient boosting (XGBRegressor; XGBoost). Feature-set definitions and graph-aligned sample availability for each cohort are provided in **Supplementary Table S2.**

### D. Missing Data and Feature Availability

Missingness was handled at the feature-set level. A session was included in a cohort-by-feature-set analysis only if it had a valid graph file, chronological age, participant/session identifier, and all covariates required by that feature set. Imaging-only models therefore retained sessions with valid graph-derived imaging inputs, whereas multimodal models additionally required the corresponding demographic, APOE, biomarker, transcriptomic, body mass index, or cardiovascular variables. Variables unavailable for an entire cohort were not imputed across cohorts. This design avoided cross-cohort imputation assumptions and ensured that each model was trained and evaluated on internally consistent feature matrices.

### E. Graph Neural Network Architecture

Brain-age prediction was implemented using a graph neural network based on graph isomorphism network layers with edge attributes (GINE), which incorporate edge attributes during message passing to jointly model regional features, network topology, and connection strength for brain-age estimation. For each participant or imaging session, the connectome graph was passed through three sequential message-passing layers. Each GINE layer used a two-layer multilayer perceptron with linear transformations and rectified linear unit activations. The hidden dimension was 64. Batch normalization and ReLU activation were applied after each GINE layer. Global mean pooling was used to aggregate node embeddings. For multimodal feature sets, the graph embedding was concatenated with the corresponding graph-level and non-imaging covariate vector. The fused representation was passed through a feed-forward regression head with two hidden linear layers, ReLU activations, dropout, and a final linear output predicting chronological age. Dropout was set to 0.35. The same architecture was used across cohorts and feature sets, with input dimensions adjusted according to the available node, edge, and graph-level features.

### F. Model Training and Cross-Validation

Models were trained separately within each cohort and feature-set configuration. Cohort-specific training, rather than pooling data across cohorts, was used to limit confounding arising from differences in image acquisition, age distributions, feature availability, and sample composition. Chronological age was used as the prediction target. Models were optimized by minimizing mean squared error using the Adam optimizer, with a learning rate and weight decay of (5×10^-4^) each. Training was performed for a maximum of 250 epochs with a batch size of 16. Early stopping was applied with a patience of 20 epochs, and the model state yielding the lowest validation mean absolute error was retained for each fold.

Primary model-performance evaluation used raw out-of-fold predictions from the cognitively normal training sample. Fivefold cross-validation was used throughout. For longitudinal cohorts, folds were grouped at the participant level to ensure that repeated sessions from the same individual were not split between training and validation sets, with age stratification applied when feasible to improve balance across folds. Single-visit cohorts were evaluated using shuffled fivefold cross-validation. Target-normalization parameters were estimated exclusively within each training fold and then applied to the corresponding held-out validation fold. Cross-validation fold composition, age distributions, fold-level performance, training-loss trajectories, and age-bias diagnostics are summarized in **Supplementary Fig. S1**.

### G. Brain-Age Gap and Age-Bias Correction

Raw brain-age gap was computed as: *BAG* = *A*^%^ − *A*, where **Â** is the predicted brain age and *A* is chronological age. Because raw BAG can retain age-dependent regression bias, corrected brain-age gap (cBAG) was derived using age-bias residualization. For fold-wise correction, the association between raw BAG and chronological age was estimated using the training data within each cross-validation fold and then applied to the corresponding held-out fold. cBAG was defined as the residual deviation after subtracting the expected age-related BAG component. For downstream validation analyses, a complementary out-of-fold global correction was used. The BAG–age association was estimated across the pooled out-of-fold predictions within each cohort, and the fitted age-dependent component was subtracted from raw BAG to obtain OOF-global cBAG. For full-cohort biological and longitudinal analyses, a final cohort-specific model was trained using all cognitively normal training observations and then applied to all prediction-ready cohort sessions. The OOF-global BAG–age regression parameters estimated from the cognitively normal OOF predictions were retained and applied without refitting to the resulting full-cohort predictions to obtain full-cohort cBAG. Raw out-of-fold predicted age was retained for primary model-performance reporting, whereas cBAG was used for biological and longitudinal validation and for calibration diagnostics in the transferability analyses. Cohort-specific cBAG centiles were obtained from the empirical within-cohort cBAG distribution, with higher centiles indicating relatively older-appearing brain age within that cohort.

### H. Model-Performance Evaluation and Baseline Comparison

Model performance was summarized by cohort and feature-set configuration using mean absolute error (MAE), root mean squared error (RMSE), coefficient of determination (R²), and Pearson correlation between predicted and chronological age. Participant-cluster bootstrap resampling with 2,000 iterations was used to estimate 95% confidence intervals while preserving within-participant dependence in longitudinal cohorts. As a conventional baseline comparison, the imaging-only GNN was compared with Ridge regression, ElasticNet regression, and XGBoost trained on the fixed-length imaging representation described above, comprising vectorized connectome edge attributes, global graph metrics, and summary statistics of node-level imaging features. Classical models were evaluated using raw out-of-fold predicted age and matched cross-validation folds where available. Paired comparisons between the GNN and classical baselines used overlapping out-of-fold predictions and two-sided participant-clustered sign-flip permutation tests with 10,000 permutations. Tests evaluated differences in absolute error and squared error.

### I. Interpretability Analysis

Model attribution was evaluated using SHAP analyses of the imaging-only model, with attributions computed using GradientExplainer. Attributions were summarized across four feature classes: global graph features, node-level features, edge-derived regional summaries, and exact connectome edges. Edge-derived regional summaries were obtained by aggregating incident exact-edge attributions for each anatomical region. Cross-cohort rankings were restricted to contributors represented in all four cohorts, and exact-edge analyses were additionally restricted to connections supported by at least 100 graph sessions across cohorts. To reduce differences in attribution scale across cohorts, mean absolute SHAP values were normalized separately within each cohort and feature class by dividing each feature’s mean absolute SHAP value by the cohort-specific sum of mean absolute SHAP values within that feature class. These within-cohort scaled values were then averaged across cohorts to generate the primary cross-cohort rankings. Raw mean absolute SHAP rankings were retained as sensitivity analyses.

A targeted anatomical analysis evaluated a prespecified set of 28 connections involving medial temporal lobe and AD–relevant regions, including the hippocampus, entorhinal cortex, parahippocampal cortex, amygdala, and selected cortical and subcortical regions. The target set was defined independently of SHAP magnitude, and the complete edge list is provided in **Supplementary Table S7**. Scaled SHAP importance for the prespecified edges was compared with the background exact-edge distribution using a one-sided Mann–Whitney U test. Empirical enrichment was also evaluated using 10,000 randomly sampled edge sets of equal size, with the observed mean and median importance compared with the random-set distributions. Global ranks and the proportion of prespecified edges falling within the top 5% of the complete exact-edge ranking were also summarized.

### J. Downstream and Longitudinal Validation

Corrected brain-age gap values were evaluated in downstream biological, clinical, and longitudinal analyses. Cross-sectional analyses tested associations between cBAG and neuroimaging-derived measures, graph-theoretic metrics, cognitive variables, biomarker features, transcriptomic principal components, and cardiometabolic variables when available. Continuous associations were quantified using Pearson correlation, with Benjamini–Hochberg false-discovery-rate correction applied within prespecified analysis families. Binary-outcome discrimination analyses evaluated APOE ε4 carrier status, cognitive impairment, and sex using area under the receiver-operating-characteristic curve (AUC). Risk-factor specificity analyses used standardized linear regression for continuous outcomes and logistic regression for binary outcomes; effects were reported as standardized β coefficients or log odds ratios per 1-SD higher cBAG, respectively, with multiple-testing correction applied within prespecified outcome families. For AD-DECODE transcriptomic analyses, preranked gene-set enrichment analysis was applied to PC17 gene loadings using the MSigDB Hallmark 2020 gene-set library, and FDR-adjusted q values were used to identify significant pathways. Longitudinal analyses were performed in ADNI and HABS-HD using repeated observations paired within participant. Change–change analyses tested associations between within-person change in cBAG and within-person change in biological endpoints using Pearson correlation, with Benjamini–Hochberg correction applied separately within each cohort across the valid endpoints tested. Longitudinal preservation of brain-age position was evaluated using baseline-to-latest-visit pairs and quantified using Pearson and Spearman correlations and ICC(2,1), with participant-level bootstrap 95% confidence intervals. Prospective change models tested whether baseline cBAG predicted subsequent endpoint change while adjusting for the baseline endpoint value, baseline age, sex, and follow-up interval when estimable. Benjamini–Hochberg correction was applied separately within cohort across the unique prospective endpoints tested.

### K. Cross-Cohort Transferability and Calibration Sensitivity

Cross-cohort transferability was evaluated using imaging-only models in a train–test grid spanning ADNI, Duke/UNC ADRC, HABS-HD, and AD-DECODE. Diagonal cells represented within-cohort out-of-fold reference performance, whereas off-diagonal cells represented external transfer without model retraining. Raw transfer performance was assessed by comparing predicted age with chronological age. Residualized corrected brain-age gap (cBAG) was additionally used to characterize remaining age dependence after bias correction. We also quantified systematic prediction offset, the frequency of implausible predicted-age values, and the number of observations with large absolute errors. Post hoc sensitivity analyses applied intercept-only and linear recalibration separately within each target cohort. Intercept-only recalibration adjusted systematic prediction offset while retaining the original prediction slope, whereas linear recalibration fit an affine mapping of transferred predicted age to chronological age, thereby adjusting both offset and scale. Recalibration parameters were estimated and evaluated within the same target cohort using chronological age; these analyses were interpreted as diagnostic calibration-sensitivity analyses rather than independent external validation. Because affine recalibration preserves participant ordering, Pearson correlation was used to assess participant-level discrimination separately from calibration-sensitive metrics such as MAE, RMSE, and R². The recalibration analyses were used to distinguish cohort-specific mismatch in prediction offset and scale from limited transportability of participant-level predictive information.

### L. Software and Reproducibility

Analyses were implemented in Python 3.9.22 using PyTorch 2.5.1 and PyTorch Geometric 2.6.1. GPU-accelerated model training was performed using CUDA-enabled PyTorch (CUDA 12.4) on one NVIDIA A100 80-GB GPU. Supporting statistical analyses used NumPy 2.0.2, pandas 2.2.3, SciPy 1.13.1, and scikit-learn 1.6.1. Conventional baseline models included Ridge regression, ElasticNet regression, and XGBoost 2.1.4. SHAP-based model attribution was implemented using SHAP 0.48.0 with GradientExplainer. Reproducibility was supported by preserving cohort-specific cross-validation fold assignments, random seeds, training histories, out-of-fold and full-cohort predictions, model-performance summaries, run manifests, quality-control outputs, execution logs, and intermediate analysis tables .

## III. RESULTS

### A. Brain-age prediction performance varied by cohort and feature-set configuration

Out-of-fold brain-age prediction performance varied across cohorts and feature-set configurations (**Fig. 2**; **Table 2**). In ADNI, errors were similar across feature sets, with the imaging-only model ranking first by MAE (MAE = 4.72 years, 95% CI: 4.21–5.26; RMSE = 5.88 years; R² = 0.159; r = 0.399). In Duke/UNC ADRC, models incorporating non-imaging variables performed better than the imaging-only model, with the full model without cardiovascular variables showing the strongest performance (MAE = 7.30 years, 95% CI: 6.46–8.18; RMSE = 9.04 years; R² = 0.363; r = 0.607). In HABS-HD, the full model ranked first and showed the strongest performance among the older-adult cohorts (MAE = 4.70 years, 95% CI: 4.25–5.16; RMSE = 6.12 years; R² = 0.431; r = 0.657). AD-DECODE was the most challenging cohort, with the imaging plus biomarkers model ranking first by MAE (MAE = 9.52 years, 95% CI: 7.66–11.60; RMSE = 13.06 years; R² = 0.255; r = 0.513), although the imaging-only model showed the best RMSE and R². Overall, prediction accuracy was more strongly influenced by cohort than by feature-set choice, and no single feature configuration was uniformly optimal across datasets.

**Figure 2.**
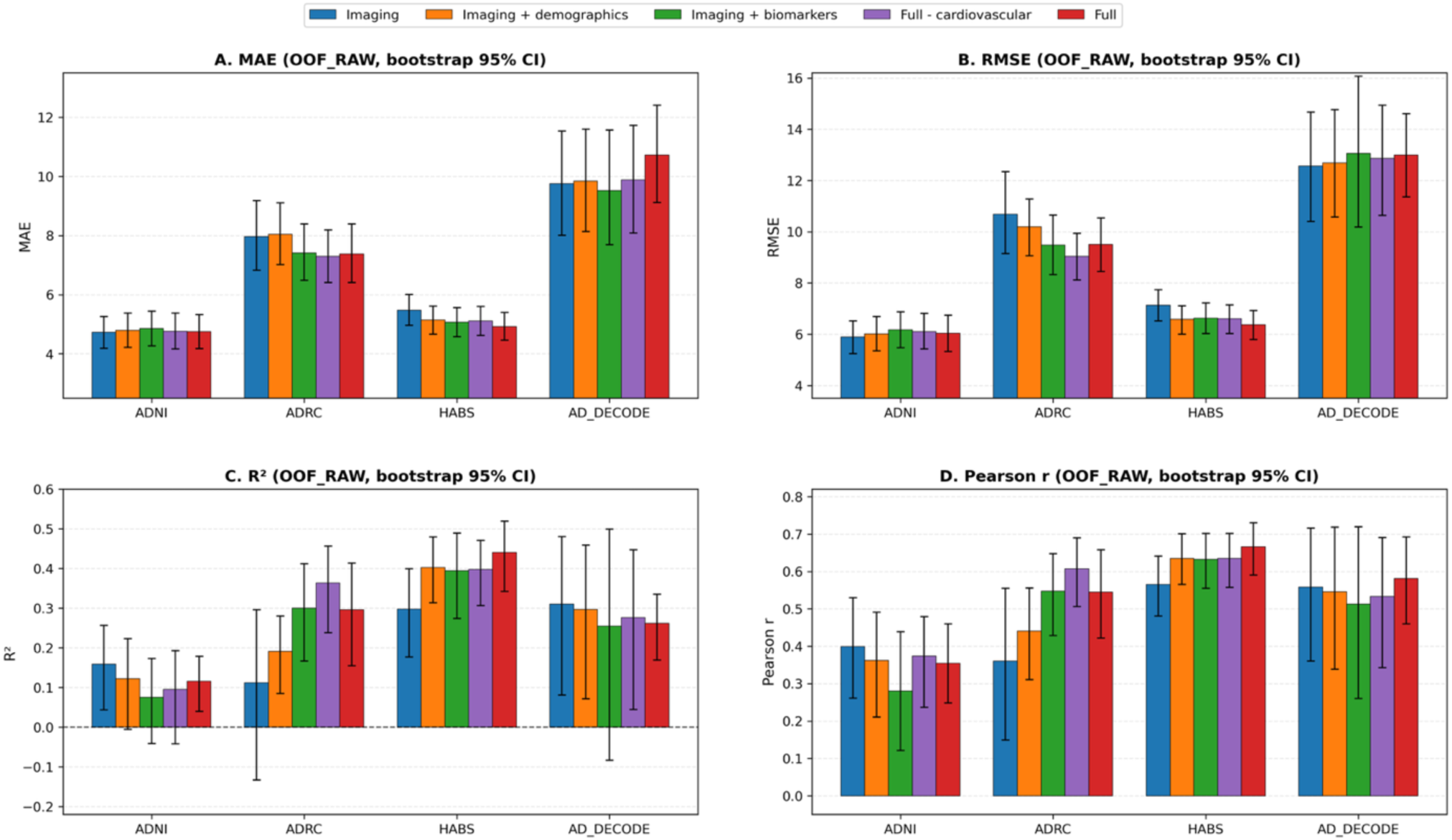
Out-of-fold raw brain-age prediction performance across cohorts and feature sets. Bar plots show out-of-fold raw predicted-age performance for five feature-set models across ADNI, Duke/UNC ADRC, HABS-HD, and AD-DECODE. Panels show **A)** mean absolute error, **B)** root mean squared error, **C)** coefficient of determination, and **D)** Pearson correlation between chronological age and predicted brain age. Error bars represent bootstrap 95% confidence intervals. Lower MAE and RMSE indicate better performance, whereas higher R² and Pearson r indicate stronger agreement between predicted and chronological age. Performance varied more strongly by cohort than by feature set, with ADNI and HABS-HD showing the lowest prediction errors and AD-DECODE showing the largest errors.

**Table 2.** Raw out-of-fold brain-age prediction performance by cohort and feature-set model. Raw out-of-fold predicted brain age was evaluated separately within each cohort and feature-set configuration using the cognitively normal training/cross-validation sample. Performance is reported as mean absolute error (MAE), root mean squared error (RMSE), coefficient of determination (R^2^), and Pearson correlation between predicted and chronological age, with participant-cluster bootstrap 95% confidence intervals. Lower MAE and RMSE indicate better performance, whereas higher (R^2^) and Pearson (r) indicate better agreement between predicted and chronological age. Models are ranked within each cohort by MAE. Full − CV denotes the full multimodal feature set excluding cardiovascular variables. Participant-level grouping was used during cross-validation for repeated-session cohorts to prevent leakage across visits.

| Rank | Model | MAE [95% CI] | RMSE [95% CI] | R<sup>2</sup> [95% CI] | Pearson r [95% CI] |
| --- | --- | --- | --- | --- | --- |
| **ADNI** |  |  |  |  |  |
| 1 | Imaging only | 4.724 [4.207, 5.261] | 5.883 [5.263, 6.507] | 0.159 [0.043, 0.250] | 0.399 [0.255, 0.520] |
| 2 | Full | 4.748 [4.201, 5.304] | 6.031 [5.340, 6.737] | 0.116 [0.041, 0.183] | 0.354 [0.244, 0.462] |
| 3 | Full – CV | 4.763 [4.214, 5.375] | 6.102 [5.428, 6.787] | 0.095 [-0.035, 0.200] | 0.374 [0.239, 0.486] |
| 4 | Imaging + demo. | 4.788 [4.255, 5.350] | 6.008 [5.338, 6.661] | 0.123 [-0.004, 0.226] | 0.362 [0.212, 0.490] |
| 5 | Imaging + biomarkers | 4.852 [4.276, 5.455] | 6.167 [5.458, 6.880] | 0.076 [-0.045, 0.168] | 0.280 [0.117, 0.437] |
| **Duke/UNC ADRC** |  |  |  |  |  |
| 1 | Full – CV | 7.297 [6.461, 8.181] | 9.039 [8.164, 9.895] | 0.363 [0.232, 0.460] | 0.607 [0.504, 0.690] |
| 2 | Full | 7.378 [6.435, 8.384] | 9.503 [8.471, 10.596] | 0.296 [0.150, 0.408] | 0.545 [0.418, 0.654] |
| 3 | Imaging + biomarkers | 7.407 [6.411, 8.399] | 9.480 [8.240, 10.683] | 0.300 [0.172, 0.411] | 0.548 [0.433, 0.650] |
| 4 | Imaging only | 7.967 [6.844, 9.116] | 10.678 [9.121, 12.229] | 0.112 [-0.140, 0.297] | 0.360 [0.138, 0.556] |
| 5 | Imaging + demo. | 8.036 [7.043, 9.070] | 10.189 [9.138, 11.249] | 0.191 [0.086, 0.277] | 0.441 [0.317, 0.549] |
| **HABS-HD** |  |  |  |  |  |
| 1 | Full | 4.697 [4.252, 5.160] | 6.119 [5.580, 6.668] | 0.431 [0.316, 0.519] | 0.657 [0.579, 0.724] |
| 2 | Full – CV | 4.776 [4.326, 5.272] | 6.255 [5.670, 6.850] | 0.405 [0.289, 0.500] | 0.639 [0.559, 0.710] |
| 3 | Imaging + biomarkers | 4.792 [4.291, 5.295] | 6.315 [5.619, 6.994] | 0.394 [0.253, 0.501] | 0.630 [0.539, 0.712] |
| 4 | Imaging + demo. | 4.975 [4.473, 5.538] | 6.552 [5.928, 7.191] | 0.348 [0.251, 0.432] | 0.597 [0.516, 0.669] |
| 5 | Imaging only | 5.143 [4.675, 5.665] | 6.579 [6.065, 7.134] | 0.342 [0.251, 0.423] | 0.586 [0.505, 0.657] |

**Table 2. Raw out-of-fold brain-age prediction performance by cohort and feature-set model.**
| AD-DECODE |  |  |  |  |  |
| --- | --- | --- | --- | --- | --- |
| 1 | Imaging + biomarkers | 9.521 [7.659, 11.599] | 13.058 [10.259, 16.027] | 0.255 [-0.049, 0.501] | 0.513 [0.274, 0.726] |
| 2 | Imaging only | 9.753 [8.043, 11.588] | 12.561 [10.379, 14.696] | 0.310 [0.087, 0.489] | 0.558 [0.365, 0.721] |
| 3 | Imaging + demo. | 9.838 [8.057, 11.683] | 12.685 [10.503, 14.957] | 0.297 [0.073, 0.466] | 0.546 [0.346, 0.720] |
| 4 | Full – CV | 9.878 [8.134, 11.746] | 12.865 [10.649, 15.002] | 0.277 [0.049, 0.458] | 0.533 [0.348, 0.697] |
| 5 | Full | 10.723 [9.097, 12.406] | 12.993 [11.357, 14.598] | 0.262 [0.165, 0.332] | 0.582 [0.457, 0.685] |

Out-of-fold raw predicted age was associated with chronological age for the imaging-only model in all cohorts (**Supplementary Fig. S2**). Prediction strength varied by cohort, with stronger associations in HABS-HD and AD-DECODE and weaker associations in ADNI and Duke/UNC ADRC. Across cohorts, fitted prediction slopes were shallower than the identity line, indicating regression-to-the-mean in raw predicted age. This pattern is expected in brain-age modeling and motivated the use of age-bias-corrected brain-age gap rather than raw BAG for downstream biological and longitudinal validation analyses. Age-dependence diagnostics confirmed that raw BAG was strongly negatively associated with chronological age across cohorts, whereas OOF-global cBAG showed near-zero residual association with age, supporting its use as the primary corrected brain-age phenotype for downstream analyses (**Supplementary Fig. S3**). As a secondary benchmark, the imaging-only GNN was compared with Ridge regression, ElasticNet regression, and XGBoost trained on a fixed-length representation of the same imaging inputs comprising vectorized connectome edge attributes, global graph metrics, and summary statistics of node-level imaging features (**Fig. 3**; **Supplementary Table S6A**). Across cohorts, the GNN was competitive with conventional models but was not uniformly superior. In ADNI, ElasticNet achieved the lowest MAE (4.715 years), closely followed by the GNN (4.724 years), whereas the GNN showed the lowest RMSE (5.883 years), highest R² (0.159), and highest Pearson correlation (r = 0.399). In Duke/UNC ADRC, all three conventional models showed lower MAE than the GNN; Ridge achieved the lowest MAE (7.448 years), whereas XGBoost showed the lowest RMSE (9.481 years), highest R² (0.300), and highest Pearson correlation (r = 0.549). In HABS-HD, XGBoost achieved the lowest MAE (5.031 years) and RMSE (6.364 years) and the highest R² (0.385) and Pearson correlation (r = 0.624), with ElasticNet and the GNN showing similar performance. In AD-DECODE, the GNN achieved the lowest MAE (9.753 years), whereas XGBoost showed slightly lower RMSE (12.421 years), higher R² (0.326), and higher Pearson correlation (r = 0.571).

**Figure 3.**
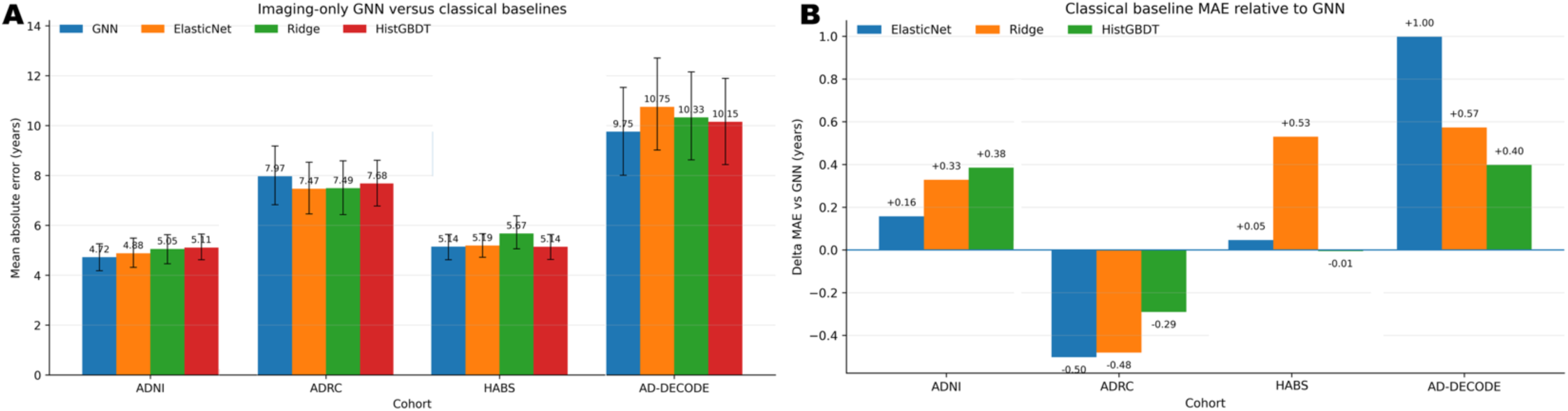
Imaging-only graph neural network performance compared with conventional regression baselines. Raw out-of-fold age-prediction performance from the imaging-only graph neural network (GNN) was compared with Ridge regression, ElasticNet regression, and XGBoost trained on a fixed-length representation of the same imaging inputs comprising vectorized connectome edge attributes, global graph metrics, and summary statistics of node-level imaging features. A, Mean absolute error (MAE) by cohort and model, with error bars indicating participant-cluster bootstrap 95% confidence intervals. B, Difference in MAE between each conventional baseline and the GNN, calculated as baseline MAE minus GNN MAE; positive values indicate lower MAE for the GNN, whereas negative values indicate lower MAE for the conventional baseline. Paired two-sided participant-clustered permutation tests used 10,000 permutations. Complete performance metrics and paired permutation results are provided in **Supplementary Tables S6A and S6B.**

Paired participant-clustered permutation tests showed broadly similar predictive performance across model classes, with isolated nominal differences in ADNI (**Supplementary Table S6B**). XGBoost showed higher absolute error than the GNN (ΔMAE = +0.322 years, baseline minus GNN; two-sided p = 0.0416), whereas Ridge showed higher squared error than the GNN (p = 0.0292). No other paired comparison reached p < 0.05. These isolated findings did not indicate consistent superiority of either the GNN or the conventional baselines across cohorts. Rather, relative performance depended on cohort and metric, supporting the interpretation that the imaging-only GNN was competitive with, but not uniformly better than, conventional models trained on the same imaging information.

Pooled analyses favored the full model without cardiovascular variables, but these pooled metrics were interpreted as supplementary because they are influenced by cohort sample size, cohort composition, and age-distribution differences. Within-cohort performance therefore remained the primary evaluation framework (**Supplementary Tables S3–S4**).

### B. Corrected brain-age gap was associated with diffusion and graph-derived network integrity

Cross-sectional biological validation evaluated associations between corrected brain-age gap (cBAG) from the imaging-only model and diffusion, volumetric, and graph-theoretic neuroimaging measures across the four cohorts (Fig. 4). The most consistent finding was a relationship between higher cBAG and reduced diffusion-derived microstructural integrity. Hippocampal fractional anisotropy (FA) was negatively associated with cBAG in ADNI (n = 316, r = −0.15, q = 0.0129), Duke/UNC ADRC (n = 204, r = −0.43, q < 10⁻⁴), and HABS-HD (n = 486, r = −0.44, q < 10⁻⁴), whereas no significant association was observed in AD-DECODE. Similarly, whole-brain FA was negatively associated with cBAG in Duke/UNC ADRC (r = −0.81, q < 10⁻⁴), HABS-HD (r = −0.60, q < 10⁻⁴), and AD-DECODE (r = −0.27, q = 0.0162), while the weaker association in ADNI did not survive false-discovery-rate correction. These findings demonstrate that older-appearing brain connectome profiles were consistently associated with lower white-matter microstructural integrity across independent cohorts.

**Figure 4.**
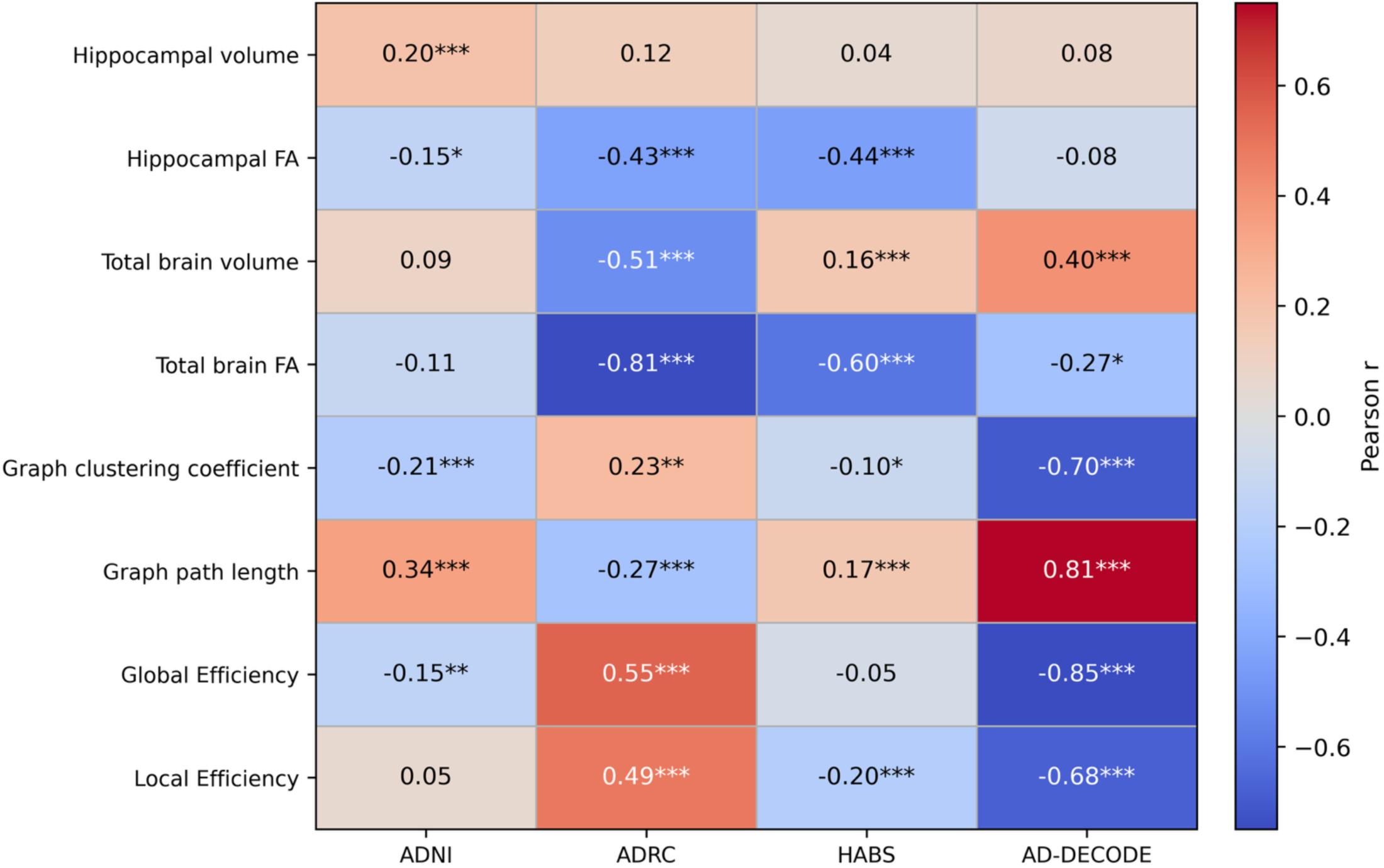
Cross sectional neuroimaging associations with imaging-only corrected brain-age gap across cohorts. Heatmap colors and cell values represent Pearson correlation coefficients between corrected brain-age gap and regional or global neuroimaging measures in ADNI, Duke/UNC ADRC, HABS, and AD-DECODE. Asterisks denote significance after Benjamini–Hochberg false-discovery-rate correction across the cohort-by-measure tests: ∗q<0.05, ∗∗q<0.01, and ∗∗∗q<0.001. Exact FDR-adjusted q-values and complete-case sample sizes are shown in each cell. Positive correlations indicate higher values of the neuroimaging measure with higher cBAG; negative correlations indicate lower values with higher cBAG.

Graph-theoretic measures also showed significant associations with cBAG, although their direction varied across datasets. Higher cBAG was associated with longer graph path length in ADNI (r = 0.34, q < 10⁻⁴), HABS-HD (r = 0.17, q < 0.001), and AD-DECODE (r = 0.81, q < 10⁻⁴), but with shorter path length in Duke/UNC ADRC (r = −0.27, q < 0.001). Likewise, graph clustering coefficient was negatively associated with cBAG in ADNI (r = −0.21, q < 0.001), HABS-HD (r = −0.10, q = 0.0367), and AD-DECODE (r = −0.70, q < 10⁻⁴), but positively associated in Duke/UNC ADRC (r = 0.23, q = 0.00137). Global and local efficiency exhibited similarly cohort-dependent patterns, with strong negative associations in AD-DECODE, positive associations in Duke/UNC ADRC, and weaker or absent effects in the remaining cohorts. These heterogeneous graph-theoretic findings likely reflect differences in network topology, acquisition protocols, preprocessing pipelines, and cohort composition rather than a single conserved pattern of connectome aging.

Volumetric measures were less reproducible than diffusion metrics. Hippocampal volume relative to total brain volume was positively associated with cBAG only in ADNI (r = 0.20, q < 0.001). Associations with total brain volume differed across cohorts, with higher cBAG corresponding to lower total brain volume in Duke/UNC ADRC (r = −0.51, q < 10⁻⁴), but higher total brain volume in HABS-HD (r = 0.16, q < 0.001) and AD-DECODE (r = 0.40, q < 0.001). Overall, these findings support the biological validity of cBAG as an imaging-derived phenotype reflecting reduced white-matter integrity. The most reproducible associations involved diffusion-derived microstructural measures: whole-brain FA was negatively associated with cBAG in Duke/UNC ADRC, HABS-HD, and AD-DECODE, whereas hippocampal FA was negatively associated with cBAG in ADNI, Duke/UNC ADRC, and HABS-HD. Volumetric and graph-theoretic associations were more heterogeneous in magnitude and direction, indicating greater cohort dependence than the diffusion-integrity signal.

### C. Cross-sectional biological validation showed strongest alignment with structural and network integrity

We next evaluated whether imaging-only corrected brain-age gap (cBAG) was associated with biologically and clinically relevant variation across cohorts. **Fig. 5(a)** summarizes the strongest cohort-specific association within each validation domain. The most consistent cross-cohort signal emerged from the imaging and network domain. Higher cBAG was associated with lower global efficiency in ADNI (r = −0.44, q = 1.77 × 10⁻¹⁴) and AD-DECODE (r = −0.85, q = 2.85 × 10⁻²⁴), and with lower total brain fractional anisotropy (FA) in Duke/UNC ADRC (r = −0.81, q = 5.63 × 10⁻⁴⁸) and HABS-HD (r = −0.60, q = 8.63 × 10⁻⁴⁸). Thus, all four cohorts showed FDR-significant associations linking higher cBAG to lower diffusion- or connectome-derived structural integrity, despite differences in the specific measure that provided the strongest association within each cohort. Molecular associations were significant but more dependent on cohort-specific data availability. In HABS-HD, higher cBAG was associated with higher plasma phosphorylated tau (pTau; r = 0.24, q = 3.59 × 10⁻⁵, n = 359). Higher cBAG was also associated with plasma Aβ40 (r = 0.53, q = 2.99 × 10⁻², n = 25), although this estimate should be interpreted cautiously because of the limited sample size.

**Figure 5.**
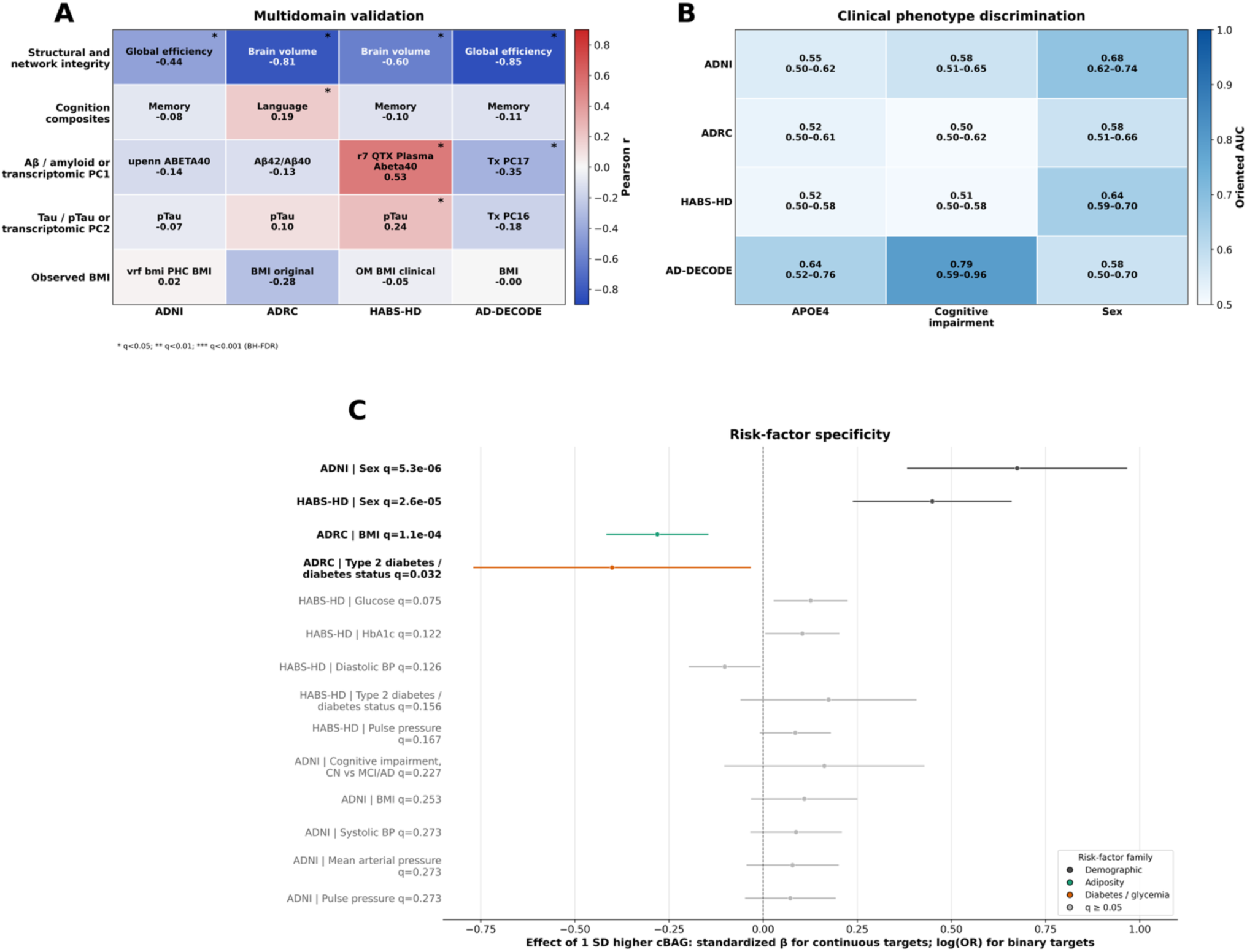
Cross-sectional biological and clinical validation of imaging-only corrected brain-age gap. Cross-sectional validation analyses tested whether corrected brain-age gap (cBAG) from the imaging-only model was associated with biological, clinical, molecular, cognitive, and risk-factor measures across ADNI, Duke/UNC ADRC, HABS-HD, and AD-DECODE. **A,** multidomain validation heatmap showing the top cohort-specific association within each validation domain. Cell colors indicate Pearson correlation coefficients, and annotations report the selected endpoint, correlation coefficient, and sample size. Asterisks indicate FDR-significant associations within the tested domain. The strongest and most consistent associations were observed in the imaging/network domain, where higher cBAG was associated with lower global efficiency in ADNI and AD-DECODE and lower total brain FA in Duke/UNC ADRC and HABS-HD. **B,** discrimination analyses for selected binary outcomes, including APOE4 carrier status, cognitive impairment, and sex. Cells show oriented AUC values with 95% confidence intervals and sample sizes. **C,** risk-factor specificity analysis showing the top-ranked demographic, adiposity, glycemic/diabetes, and vascular associations. Points show the effect of 1 SD higher cBAG, with horizontal bars indicating 95% confidence intervals. Effects are standardized β coefficients for continuous targets and log odds ratios for binary targets. FDR-significant rows are shown in family color; non-significant rows are shown in gray. Overall, imaging-only cBAG aligned most strongly with imaging/network integrity, whereas non-imaging risk-factor associations were smaller and cohort-specific.

In AD-DECODE, transcriptomic principal components were available instead of fluid biomarkers. Higher cBAG was associated with lower PC17 values (r = −0.35, q = 3.72 × 10⁻², n = 77), placing higher-cBAG individuals toward the negative-loading side of this latent transcriptomic component. Preranked gene-set enrichment analysis of the PC17 gene loadings identified three Hallmark pathways that survived FDR correction. Positive PC17 loadings, corresponding to the lower-cBAG direction, were enriched for heme metabolism (NES = 2.56, q = 0.001) and pancreas beta-cell signatures (NES = 1.84, q = 0.008). Negative PC17 loadings, corresponding to the higher-cBAG direction, were enriched for interferon-alpha response (NES = −1.76, q = 0.014; **Supplementary Table S5**). These results suggest that the transcriptomic correlate of higher cBAG in AD-DECODE was aligned with an interferon-related immune-response axis, whereas lower cBAG was associated with heme-metabolic and beta-cell-related signatures. This pathway-level interpretation remains exploratory because PC17 is a latent transcriptomic component, its biological direction depends on the structure of its gene loadings, and independent molecular replication is required.

Cognitive associations were weaker than the imaging and network associations. Associations with the selected cognitive composites were generally modest and did not show a reproducible FDR-significant pattern across cohorts. BMI likewise showed limited and cohort-specific evidence, with the clearest association observed in Duke/UNC ADRC, where higher cBAG was associated with lower BMI.

Overall, cross-sectional validation indicated that cBAG aligned most consistently with reduced structural and network integrity across cohorts. Molecular correlates provided additional evidence of disease-relevant biological variation, including plasma pTau and an interferon-related transcriptomic axis, but were constrained by cohort-specific biomarker availability. In contrast, cognitive and BMI associations were weaker and less reproducible.

**Fig. 5(b)** evaluated whether cBAG discriminated APOE4 carrier status, cognitive impairment, and sex. Overall, discrimination was modest. Oriented AUCs for APOE4 carrier status ranged from 0.52 to 0.64, cognitive-impairment AUCs ranged from near chance to 0.79, and sex AUCs ranged from 0.58 to 0.68. The strongest cognitive-impairment discrimination was observed in AD-DECODE, but the small sample size and cohort-specific context require caution. These findings indicate that imaging-only cBAG did not function as a strong diagnostic, genetic-risk, or sex discriminator. Risk-factor specificity analyses further supported this interpretation (**Fig. 5(c)**). After FDR correction, higher cBAG was associated with male sex in ADNI and HABS-HD, and with lower BMI and lower odds of type 2 diabetes/diabetes status in Duke/UNC ADRC. Other glycemic, vascular, and clinical-status associations were weaker and did not survive FDR correction. Strict APOE4 carriage and cognitive-impairment status were not consistently associated with cBAG. Overall, **Fig. 5** shows that imaging-only cBAG aligned most strongly with continuous imaging/network integrity, whereas molecular, cognitive, demographic, and cardiometabolic associations were weaker, exploratory, or cohort-specific.

### D. Longitudinal preservation of cBAG and prediction of subsequent imaging deterioration

Longitudinal validation was performed in ADNI and HABS-HD using cBAG derived from the imaging-only model. We evaluated whether within-person change in cBAG tracked contemporaneous biological and clinical change, whether participants maintained their relative cBAG position across visits, and whether baseline cBAG predicted subsequent imaging and network deterioration. In longitudinal change–change analyses, Benjamini–Hochberg FDR correction was applied separately within each cohort across all valid tested endpoints. In ADNI, greater within-person increases in cBAG were associated with greater decreases in diastolic blood pressure (n = 136, r = −0.405, q < 0.001). In HABS-HD, greater increases in cBAG were associated with greater increases in graph path length (n = 168, r = 0.270, q = 0.0057). No other cohort-specific change–change association survived FDR correction. Complete change–change results, including nominal associations, are provided in **Supplementary Fig. S4.** We next assessed whether cBAG preserved each participant’s relative brain-aging position across repeated visits. In ADNI, 136 participants had baseline-to-latest-visit measurements separated by a median interval of approximately 4 years. Baseline and follow-up cBAG showed moderate preservation (Spearman’s ρ = 0.64, Pearson’s r = 0.67, ICC(2,1) = 0.67, 95% CI: 0.53–0.78), with a median absolute within-person cBAG change of 1.4 years. Preservation was stronger in HABS-HD, where 168 participants were followed over a median interval of approximately 2 years (ρ = 0.80, r = 0.82, ICC(2,1) = 0.81, 95% CI: 0.75–0.86), with a median absolute within-person change of 1.9 years. Cohort-specific cBAG centiles showed similar preservation, with ICC(2,1) = 0.64 (95% CI: 0.53–0.75) in ADNI and ICC(2,1) = 0.79 (95% CI: 0.73–0.85) in HABS-HD. The median absolute centile change was 13.2 points in ADNI and 10.4 points in HABS-HD; 43% and 47% of participants, respectively, remained within 10 centile points of baseline, while 41% and 45% remained within the same centile quintile. These findings indicate that cBAG captured a partially stable, participant-specific position within the cohort brain-aging distribution. Finally, prospective models evaluated whether baseline cBAG predicted subsequent imaging and network change after adjustment for the baseline outcome value, baseline age, sex, and follow-up interval where estimable. Benjamini–Hochberg correction was applied separately across 15 unique prospective endpoints in ADNI and 10 endpoints in HABS-HD. In ADNI, higher baseline cBAG predicted subsequent decreases in hippocampal fractional anisotropy (FA; β = −0.285, 95% CI: −0.428 to −0.142, q = 0.0016), local efficiency (β = −0.242, 95% CI: −0.365 to −0.118, q = 0.0016), whole-brain FA (β = −0.240, 95% CI: −0.384 to −0.096, q = 0.0074), and clustering coefficient (β = −0.187, 95% CI: −0.323 to −0.050, q = 0.0264), together with an increase in hippocampal radial diffusivity (β = 0.223, 95% CI: 0.051–0.395, q = 0.0332). The directions of these associations consistently linked higher baseline cBAG to subsequent microstructural and network deterioration. No prospective imaging or network association survived FDR correction in HABS-HD. **Supplementary Fig. S5A** presents annualized longitudinal change metrics accounting for differences in follow-up duration. **Supplementary Fig. S5B** presents corrected predicted-age and predicted-age-centile preservation. Median annualized absolute cBAG change was 0.36 years per year in ADNI and 0.97 years per year in HABS-HD; the corresponding median annualized absolute centile changes were 3.31 and 5.21 centile points per year, respectively. Together, these findings show that imaging-only cBAG was moderately to strongly preserved within individuals while remaining sensitive to selected longitudinal biological changes. Importantly, higher baseline cBAG prospectively identified ADNI participants who subsequently exhibited greater diffusion-microstructural and network deterioration, providing longitudinal evidence that cBAG captures biologically meaningful variation. These longitudinal findings are summarized in **Fig. 6**.

**Figure 6.**
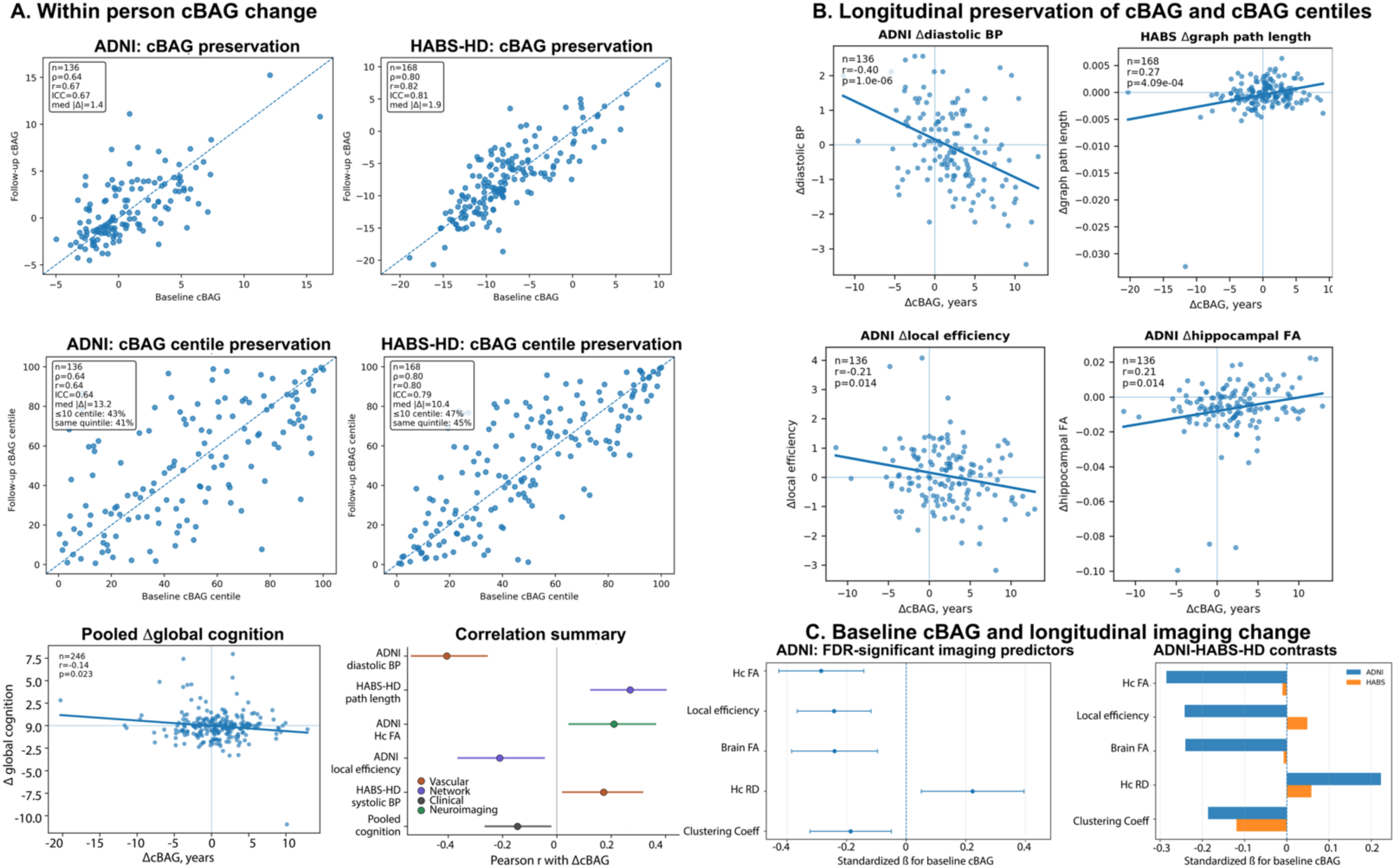
Longitudinal cBAG biological change and preservation of brain-age position. Higher baseline cBAG predicted subsequent deterioration of diffusion and network integrity in ADNI. Longitudinal analyses were performed in ADNI and HABS-HD using full-cohort cBAG estimates from the imaging-only model. A, Longitudinal associations between within-person change in corrected brain-age gap (ΔcBAG) and within-person change in selected biological endpoints. Panel A includes cohort-specific and pooled analyses for the strongest longitudinal associations identified in the full-cohort analysis. Scatterplots show ΔcBAG versus endpoint change, and the summary subpanel shows correlation coefficients with horizontal error bars indicating 95% confidence intervals. B, Longitudinal preservation of cBAG and cBAG centiles. Baseline-to-latest-visit pairs were evaluated in 136 ADNI participants over a median interval of 4 years and 168 HABS-HD participants over a median interval of 2 years. Scatterplots compare baseline and follow-up cBAG values or cohort-specific cBAG centiles; dashed lines indicate identity. Insets report the number of longitudinal pairs, Spearman’s ρ, Pearson’s r, ICC(2,1) with nonparametric bootstrap 95% confidence intervals, median absolute difference, and, for centiles, the percentage of pairs remaining within 10 centile points and within the same quintile. cBAG showed moderate longitudinal preservation in ADNI (ρ = 0.64, r = 0.67, ICC = 0.67 [0.53, 0.78]) and stronger preservation in HABS-HD (ρ = 0.80, r = 0.82, ICC = 0.81 [0.75, 0.86]). cBAG centiles were also preserved over time, with ICC = 0.64 [0.53, 0.75] in ADNI and ICC = 0.79 [0.73, 0.85] in HABS-HD. Median absolute cBAG-centile change was 13.2 centile points in ADNI and 10.4 centile points in HABS-HD. Together, these analyses show that cBAG demonstrates moderate-to-good longitudinal stability while still capturing meaningful within-person biological change.

### E. Cross-cohort SHAP analysis identifies distributed network contributors to predicted brain age

SHAP analysis identified features that contributed most strongly to predicted brain age in the imaging-only model. SHAP values were summarized across ADNI, Duke/UNC ADRC, HABS-HD, and AD-DECODE within four feature classes: global graph measures, node-level features, edge-derived regional summaries, and exact connectome edges. To reduce dominance by cohorts with larger absolute attribution scales, the primary cross-cohort analysis used mean absolute SHAP values normalized within cohort and feature class before averaging across cohorts. Contributors shown in **Fig. 7** were restricted to features present in all four cohorts; exact-edge analyses were additionally restricted to connections supported by at least 100 graph sessions across cohorts.

**Figure 7.**
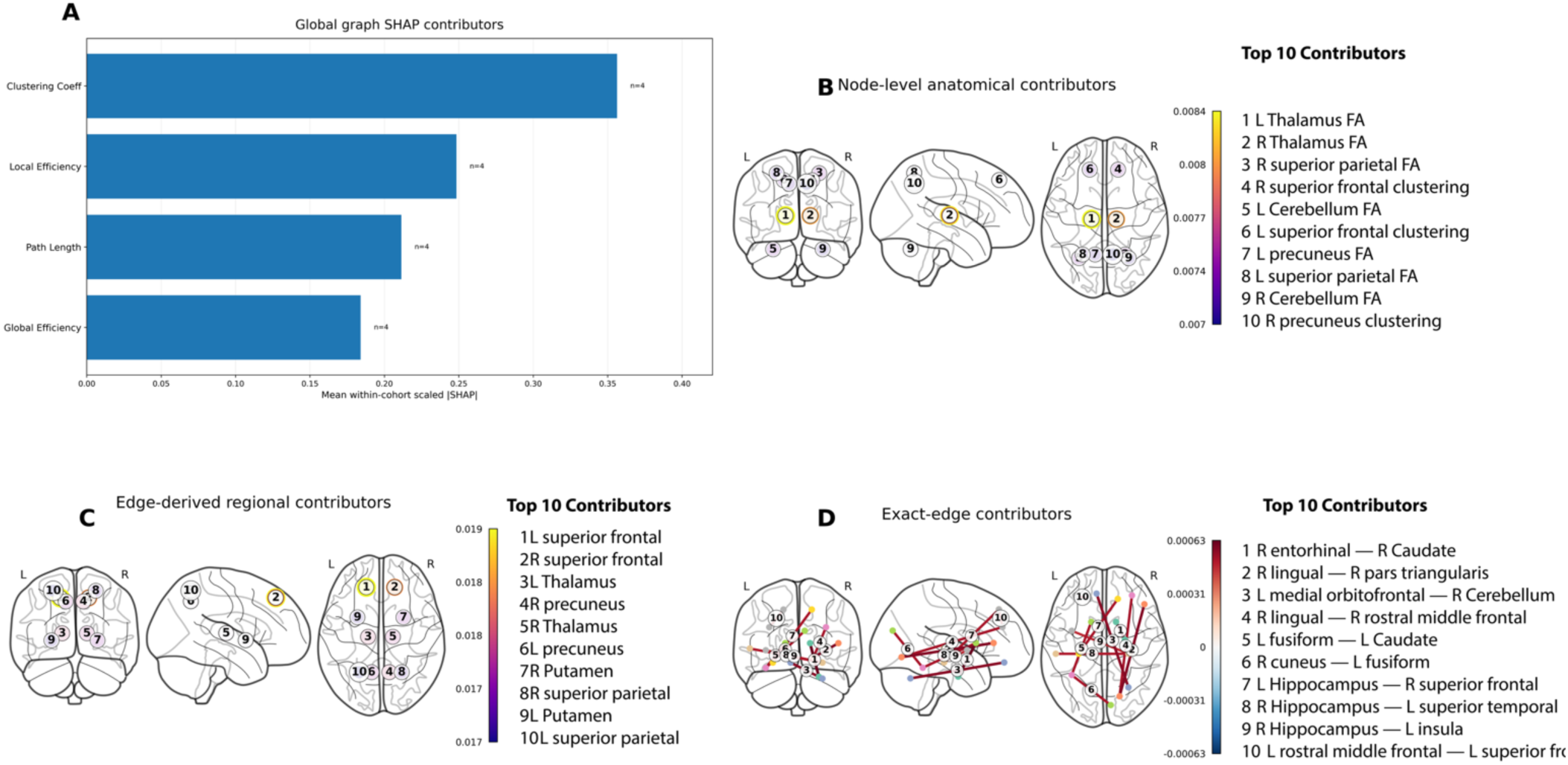
Cross-cohort multiscale SHAP contributors to predicted brain age in the imaging-only model. SHAP attributions were summarized across ADNI, Duke/UNC ADRC, HABS-HD, and AD-DECODE for features available in all four cohorts. Mean absolute SHAP values were normalized within cohort and feature class before cross-cohort averaging to reduce differences in attribution scale across cohorts. A, Global graph contributors ranked by mean within-cohort scaled absolute SHAP value; annotations indicate the number of cohorts contributing to each summary. B, Top 10 node-level contributors, including regional fractional anisotropy and nodewise graph features. Numbered brain labels correspond to the ranked list, and color indicates mean scaled absolute SHAP value. C, Top 10 edge-derived regional contributors obtained by aggregating incident edge-level SHAP values for each region. D, Top 10 exact-edge contributors, restricted to edges present in all four cohorts and supported by at least 100 graph sessions across cohorts. Line color indicates the signed cross-cohort mean SHAP value, whereas node colors distinguish anatomical endpoints. SHAP values are interpreted as descriptive model attributions rather than inferential coefficients or causal effects.

At the global graph level, clustering coefficient was the highest-ranked contributor, followed by local efficiency, path length, and global efficiency [**Fig. 7(a)**]. This ordering was stable in both raw and within-cohort scaled analyses, indicating consistent contributions of global network topology, particularly clustering- and efficiency-related properties, to predicted brain age across cohorts.

Node-level SHAP contributors were distributed across subcortical, association cortical, and cerebellar regions [**Fig. 7(b)**]. The highest-ranked scaled contributors included bilateral thalamic fractional anisotropy, superior parietal fractional anisotropy, superior frontal nodewise clustering, cerebellar fractional anisotropy, precuneus features, caudate volume, and putamen fractional anisotropy. These findings indicate that the imaging-only model relied on distributed regional diffusion, volumetric, and graph-derived information rather than on a single dominant anatomical feature. Edge-derived regional SHAP summaries showed a similarly distributed pattern [**Fig. 7(c)**]. Regions with the largest aggregated incident edge attributions included superior frontal cortex, thalamus, precuneus, putamen, superior parietal cortex, hippocampus, caudate, cerebellum, and insula. These summaries provide an intermediate level of interpretation between node-specific features and individual structural connections, highlighting fronto-subcortical, parietal–precuneus, cerebellar, insular, and medial temporal network components.

Raw-versus-scaled sensitivity analyses supported the use of within-cohort scaled SHAP values for the primary cross-cohort interpretation (**Supplementary Fig. S6**). Global graph-feature rankings were stable across raw and scaled analyses, whereas node-level, edge-derived regional, and exact-edge rankings showed partial reordering after normalization. Raw SHAP values therefore provide an absolute-magnitude reference, whereas within-cohort scaling provides a more balanced summary of attribution patterns across heterogeneous cohorts.

Exact-edge SHAP analysis provided greater anatomical specificity [**Fig. 7(d)**]. Among the highest-ranked within-cohort scaled connections were right entorhinal cortex–right caudate, right lingual cortex–right pars triangularis, left medial orbitofrontal cortex–right cerebellar cortex, right lingual cortex–right rostral middle frontal cortex, left fusiform cortex–left caudate, right cuneus–left fusiform cortex, left hippocampus–right superior frontal cortex, right hippocampus–left superior temporal cortex, and right hippocampus–left insula. These contributors further support a distributed network pattern involving temporal–frontal–striatal, cerebellar, occipital–temporal, and medial temporal circuitry. Extended anatomical visualizations of the top 30 node-level, edge-derived regional, and exact-edge contributors are provided in **Supplementary Fig. S7(a)–(c),** respectively.

Together, these analyses indicate that predicted brain age in the imaging-only model was supported by distributed connectome features showing cross-cohort consistency across multiple anatomical scales. Global graph topology contributed consistently, whereas regional and edge-level attributions emphasized thalamic, striatal, superior frontal, superior parietal, precuneus, cerebellar, insular, hippocampal, and entorhinal circuitry. SHAP values were interpreted as descriptive model attributions rather than inferential coefficients or evidence of causal anatomical effects.

Because several highly ranked exact-edge contributors involved medial temporal and Alzheimer’s disease– relevant regions, we performed a targeted descriptive analysis of a prespecified set of connections involving hippocampal, entorhinal, parahippocampal, amygdalar, and related anatomical regions. The targeted anatomical visualization is shown in Supplementary **Fig. S7(d)**, the complete prespecified edge set is reported in **Supplementary Table S7**, and enrichment relative to the full exact-edge distribution is summarized in **Supplementary Table S8**. Among 1,816 ranked exact edges, all 28 prespecified MTL/AD-relevant connections were represented. Mean within-cohort scaled absolute SHAP importance was higher for the MTL/AD-relevant set than for background edges (1.91 × 10⁻⁴ versus 1.53 × 10⁻⁴; one-sided Mann–Whitney U = 37,484, p < 0.0001). Median importance was likewise higher for MTL/AD-relevant edges (1.90 × 10⁻⁴ versus 1.55 × 10⁻⁴). In permutation testing using 10,000 equal-size random edge sets, both the observed mean and median importance exceeded the corresponding random-set distributions (empirical p < 0.0001 for both). The prespecified edges had a median global rank of 296 (range, 7–1365), and 12 of 28 (42.9%) were within the top 5% of the complete exact-edge ranking. These findings indicate that the prespecified MTL/AD-relevant edge set showed greater model attribution than the background exact-edge distribution. The targeted analysis therefore provides complementary anatomical context for the unbiased exact-edge rankings in **Fig. 7(d)**. SHAP values were interpreted as descriptive model attributions rather than inferential coefficients or evidence of causal anatomical effects.

### F. Cross-cohort transferability and post-hoc recalibration

Exploratory cross-cohort transferability analyses showed substantial sensitivity to cohort shift (**Supplementary Fig. S8; Supplementary Tables S9–S10**). Imaging-only models were trained in one cohort and evaluated in each of the remaining cohorts, with diagonal cells representing within-cohort out-of-fold reference performance and off-diagonal cells representing external transfer without retraining. Within-cohort raw MAE ranged from 4.72 years in ADNI to 9.75 years in AD-DECODE, whereas several external-transfer cells showed substantially larger errors. The most extreme failures occurred when AD-DECODE was used as the training cohort and applied to ADNI, Duke/UNC ADRC, or HABS-HD, with raw MAE values of 80.57, 55.13, and 77.83 years, respectively. Mean absolute cBAG and cBAG-age slope diagnostics similarly indicated cohort-specific calibration instability under external transfer. These analyses were interpreted as deployment stress tests rather than primary model-performance estimates and support cohort-aware validation before external clinical use.

To determine whether poor external transfer primarily reflected systematic calibration mismatch or loss of participant-level predictive information, we next performed post-hoc recalibration sensitivity analyses across all 16 train–test cohort combinations (**Supplementary Fig. S9; Supplementary Table S11**). Raw predictions were compared with intercept-only recalibration and linear recalibration using target-cohort chronological age. Diagonal cells represented within-cohort out-of-fold performance, whereas off-diagonal cells represented external transfer. Within-cohort performance changed minimally following recalibration, indicating that cohort-specific models were already well calibrated. Across the four diagonal cells, median MAE remained essentially unchanged (6.55, 6.56, and 6.66 years for raw, intercept-only, and linear recalibration, respectively), while median R² remained stable at approximately 0.24 under all three conditions.

In contrast, recalibration substantially improved external-transfer performance. Across the 12 off-diagonal cells, median MAE decreased from 17.39 years for raw predictions to 9.75 years after intercept-only recalibration and 8.22 years after linear recalibration. Median RMSE decreased from 22.68 to 13.19 and 9.98 years, respectively. Median R² increased from −6.78 before recalibration to −0.13 following intercept-only recalibration and 0.03 after linear recalibration. Whereas all raw external-transfer R² values were negative, all became non-negative following linear recalibration (range, 0.001–0.264). Recalibration also markedly reduced prediction failures. Across external-transfer evaluations, raw predictions included 761 predicted ages below 0 years, 7 above 120 years, and 927 predictions with absolute errors exceeding 50 years. These counts decreased to 0, 6, and 7, respectively, after intercept-only recalibration and were completely eliminated following linear recalibration. Despite these improvements in absolute agreement, recalibration did not alter participant-level discrimination. Median Pearson correlation across external-transfer cells remained 0.171 under raw, intercept-only, and linear recalibration. Thus, recalibration corrected systematic differences in prediction offset and scale without improving the relative ordering of individuals within the target cohort. Together, these findings indicate that poor raw cross-cohort performance reflected both systematic calibration mismatch and limited transportability of participant-level predictive information.

## IV. DISCUSSION

This study presents a calibration-aware and hierarchically interpretable framework for diffusion-connectome brain-age modeling across four heterogeneous aging and Alzheimer’s disease–related cohorts. This framework addresses several challenges that continue to limit the interpretation and translation of brain-age models: age-dependent prediction bias, incomplete biological and longitudinal validation, restricted model interpretability, and uncertain performance under cohort shift. Across 1,093 diffusion MRI/connectome sessions from 789 participants, prediction performance varied more strongly by cohort than by feature configuration; the imaging-only graph neural network was competitive with conventional regression models but was not uniformly superior. This cohort dependence likely reflects both sampling variability and systematic differences in acquisition protocols, diffusion data quality, tractography, demographic composition, and clinical heterogeneity. Cohorts with restricted age ranges or substantial disease-related connectomic variability may also provide a less well-defined normative aging trajectory, reducing chronological-age prediction accuracy. Importantly, poorer age prediction does not necessarily imply that BAG is biologically less informative, because deviations from the normative age trajectory may themselves reflect disease- or risk-related alterations. Corrected brain-age gap (cBAG) aligned most consistently with diffusion-derived microstructural and structural-network integrity; longitudinal analyses demonstrated preservation of individual brain-aging position and prediction of subsequent imaging deterioration in ADNI; and external transfer revealed distinct contributions of calibration mismatch and limited participant-level discrimination. These findings move diffusion-connectome brain-age modeling beyond an accuracy-centered prediction task toward a more rigorous evaluation of imaging-biomarker validity, interpretability, and transportability (Cole and Franke 2017) (Gaser, Kalc, and Cole 2024) (Collins et al. 2024).

Brain-age research has traditionally emphasized prediction error and the association between predicted and chronological age. Although these measures remain necessary, they do not establish that BAG is biologically meaningful, longitudinally informative, or suitable for application outside the development population. A central contribution of the present work is the explicit separation of predictive performance from biological interpretation. Regression toward the sample mean can cause raw BAG to remain associated with chronological age and thereby confound relationships with age-related phenotypes. Here, age-bias correction was estimated without using held-out observations, whereas raw out-of-fold predicted age remained the basis for primary performance assessment. This distinction preserves an unbiased estimate of predictive accuracy while providing a less age-dependent cBAG phenotype for downstream validation (Butler et al. 2021) (de Lange et al. 2022). Predicted age and corrected BAG should be regarded as related but distinct model outputs: the former evaluates age prediction, whereas the latter supports biological and longitudinal interpretation.

Prediction performance depended more strongly on cohort characteristics than on the inclusion of additional feature blocks. Multimodal inputs improved performance in some settings, particularly Duke/UNC ADRC and HABS-HD, but no feature configuration was consistently optimal across all four cohorts. This likely reflects differences in age range, sample size, diagnostic composition, acquisition and preprocessing, and the availability and measurement quality of demographic, cardiovascular, molecular, and transcriptomic variables. The finding cautions against the assumption that adding modalities necessarily produces a more robust brain-age model. Non-imaging variables may improve within-cohort prediction when they capture cohort-specific age-related variation, yet reduce comparability or portability when definitions, assay platforms, missingness patterns, and covariate distributions differ across studies. The imaging-only model therefore provided the most consistent reference for cross-cohort evaluation and the basis for interpreting brain age as an imaging-derived phenotype. The imaging-only GNN did not significantly outperform ridge regression, elastic net, or gradient boosting trained on matched vectorized connectome features. This result clarifies the contribution of graph learning. Structural connectomes are intrinsically relational: regional properties, connection attributes, and global network organization are interdependent. The GINE architecture preserved this structure and incorporated edge attributes during message passing, enabling information to be represented across node, edge, regional, and graph levels. Conventional vectorized approaches remained strong predictors, but they do not naturally preserve this multiscale network hierarchy. The present findings therefore support a more measured rationale for graph learning in medical imaging: graph architectures are most valuable when their inductive structure and interpretability align with the biological organization of the data, rather than when model complexity is assumed to guarantee superior prediction (Kawahara et al. 2017) (Cai, Gao, and Liu 2022) (Gao et al. 2023) (Kazi et al. 2025).

The strongest evidence for biological coherence was the association of higher cBAG with reduced diffusion-derived microstructural and structural-network integrity. Across cohorts, the leading imaging/network associations involved either total-brain fractional anisotropy or global efficiency, while hippocampal FA was negatively associated with cBAG in three of the four datasets. Because the imaging-only model was derived from diffusion-connectome and regional imaging features, these associations should be interpreted primarily as evidence that cBAG preserved biologically meaningful variation in tissue microstructure and network organization, rather than as fully independent validation. This pattern is biologically plausible because diffusion MRI is sensitive to age-related differences in axonal organization, myelination, and extracellular water, whereas connectome metrics summarize the consequences of these alterations for distributed network organization (Cole 2020) (Li et al. 2020; Beck et al. 2021). The cross-cohort consistency of the diffusion signal suggests that cBAG reflected a broad structural aging phenotype rather than simply gross anatomical volume or residual chronological-age effects. The persistence of cohort effects in imaging-only models suggests that study-specific differences in diffusion acquisition, preprocessing, and connectome construction may contribute to the observed domain shift. Harmonization may reduce this component of between-cohort variability, but the present results also indicate that calibration mismatch alone does not explain the limited external transportability. In particular, target-cohort recalibration substantially reduced absolute prediction error without improving participant-level age ranking. Thus, harmonization should be evaluated as one component of a broader multicohort strategy, alongside domain adaptation, broader multisite training, and explicit modeling of population heterogeneity.

The absence of a significant hippocampal FA association in AD-DECODE may partly reflect its more cognitively preserved composition, with fewer cognitively impaired participants than the other cohorts and consequently a potentially narrower range of hippocampal microstructural injury. This interpretation remains tentative, however, because differences in age distribution, sample size, acquisition, and diffusion-processing characteristics may also contribute to the cohort-specific result.

Graph-theoretic and volumetric associations were more heterogeneous in direction. Such variability may reflect genuine differences in population biology, but it also underscores the sensitivity of graph measures to parcellation, tractography, edge definition, weighting, density, thresholding, and preprocessing. Independently constructed connectomes may therefore assign different biological meaning to nominally identical measures such as clustering or efficiency. Regional and whole-brain diffusion measures may provide a more stable cross-cohort anchor for cBAG, whereas derived graph-topological measures may be more dependent on acquisition and processing context. This issue is particularly relevant for multicenter connectome studies, in which harmonization of images alone may not fully harmonize the resulting network representations.

The multidomain analyses further defined what cBAG represented. Associations with cognition, APOE4 carriage, cognitive impairment, sex, adiposity, diabetes, and vascular measures were generally weaker and less reproducible than associations with imaging and network integrity. Molecular findings involving phosphorylated tau in HABS-HD and transcriptomic PC17 in AD-DECODE were biologically suggestive but cohort specific. The present evidence therefore supports cBAG primarily as an imaging-derived measure of structural brain aging rather than as a direct diagnostic classifier, genetic-risk marker, or general index of clinical severity. At the same time, these molecular findings provide testable hypotheses regarding relationships between structural brain aging, tau-related pathological processes, and immune or metabolic pathways that can be examined in cohorts with harmonized biomarker panels.

Longitudinal validation provided an important extension beyond the predominantly cross-sectional brain-age literature. cBAG showed moderate preservation in ADNI and stronger preservation in HABS-HD, with ICC values of 0.67 and 0.81, respectively. Cohort-specific cBAG centiles were also preserved, indicating that participants tended to maintain their relative position within the brain-aging distribution. Yet cBAG was not invariant: higher baseline values predicted subsequent deterioration in hippocampal and whole-brain FA, hippocampal radial diffusivity, clustering coefficient, and local efficiency in ADNI. This combination is consistent with desirable properties of an imaging biomarker. A highly unstable measure would not reliably distinguish individuals, whereas a completely fixed measure would be insensitive to progression. The findings therefore suggest that cBAG captured a partially stable participant-specific phenotype while retaining sensitivity to subsequent structural deterioration. The stronger prospective signal in ADNI may reflect its older risk profile, longer follow-up, greater neurodegenerative burden, or differences in endpoint availability, underscoring that longitudinal validity may depend on cohort composition and observation interval.

The hierarchical SHAP analysis addresses another limitation of the field: model interpretation is often reduced to a single feature-ranking table. Brain networks are organized across multiple spatial and topological scales, and feature importance at one level cannot reveal whether predictions are driven by global topology, regional tissue properties, the connectivity of specific regions, or individual pathways. Here, attributions were examined across global graph measures, node-level features, edge-derived regional summaries, and exact structural connections. The resulting pattern was distributed, involving clustering and efficiency-related measures together with thalamic, striatal, frontal, parietal, precuneus, cerebellar, insular, hippocampal, and entorhinal circuitry. Within the a priori selected medial temporal and Alzheimer’s disease–relevant connection set, high SHAP attribution was concentrated in a restricted subset of edges rather than uniformly distributed across the full anatomical panel. This pattern supports anatomical specificity within a hypothesis-driven analysis, but it should not be interpreted as independent discovery of a medial temporal or AD circuit. SHAP values describe attribution within the fitted model rather than causal neurobiology, and future studies should assess attribution stability across resampling, model retraining, perturbation analyses, and independent datasets (Lundberg and Lee 2017) (Tjoa and Guan 2020).

The external transfer analyses revealed the principal barrier to overcome when aiming for broader application. Several off-diagonal train–test combinations produced large errors and unstable age-gap estimates. Post-hoc linear recalibration reduced median external MAE from 17.39 to 8.22 years and eliminated extreme prediction failures, but median Pearson correlation remained 0.171. External failure reflected two distinct components: calibration mismatch and loss of participant-level discrimination. Affine recalibration corrected prediction offset and scale but could not recover individual-level information that had failed to transfer. Consequently, calibration-sensitive measures such as MAE, RMSE, and *R*^2^, and ranking-oriented measures such as correlation, describe complementary dimensions of transportability and should be reported together (Van Calster et al. 2016) (Van Calster et al. 2019).

Because recalibration used chronological-age information from the target cohort, it was a diagnostic sensitivity analysis rather than deployable external validation. In a prospective setting, calibration parameters would need to be estimated in an independent calibration subset and tested in unseen participants. Even then, recalibration would not restore lost discrimination. The modest external correlations indicate that differences in age distribution, acquisition, preprocessing, cohort composition, and connectome representation affected more than prediction offset alone. Harmonization, domain adaptation, pooled or federated multicenter training, and calibration-transfer methods will therefore be necessary to improve external generalization. This interpretation is consistent with contemporary guidance emphasizing that prediction models should be evaluated not only for discrimination, but also for calibration, transparency, reproducibility, and performance in the intended use setting (Sullivan et al. 2015) (Collins et al. 2024).

Several limitations need to be acknowledged. Connectomes were generated independently within each cohort and were not harmonized before modeling, likely contributing to heterogeneous graph associations and transfer failure. Cohort-specific training reduced the mixing of incompatible feature distributions but limited effective sample size and did not evaluate pooled harmonized models. Non-imaging feature blocks differed across cohorts, limiting direct multimodal comparison and molecular replication. Longitudinal validation was available only in ADNI and HABS-HD, and prospective imaging associations survived correction only in ADNI. Some imaging validation measures were also related to information available to the imaging-only model; these associations therefore support biological coherence more strongly than fully independent validation. Finally, SHAP attribution remains descriptive and requires stability assessment and external replication.

In summary, this work shifts diffusion-connectome brain-age modeling from an accuracy-centered prediction task toward a more rigorously evaluated imaging biomarker. By integrating leakage-safe age-bias correction, hierarchical explainability, biological and longitudinal validation, and systematic external transfer testing, the framework provides a principled approach for developing and evaluating brain-age models from structural connectomes. The principal contribution is not a claim of uniform predictive superiority for graph neural networks, but a multi-cohort framework for determining whether diffusion-connectome brain-age models are adequately corrected, interpretable, biologically supported, and robust to cohort shift. Further advances in connectome harmonization, domain adaptation, and external calibration will be required before broader multicenter application, but the present work provides a foundation for evaluating the incremental value of diffusion-connectome brain age in clinical research and population neuroscience.

## Acknowledgements

We are grateful to Dr. Hussain Yassine and Dr. Meredith Braskie for helpful discussions. Data collection and sharing for the Alzheimer’s Disease Neuroimaging Initiative (ADNI) is funded by the National Institute on Aging (National Institutes of Health Grant U19AG024904). The grantee organization is the Northern California Institute for Research and Education. In the past, ADNI has also received funding from the National Institute of Biomedical Imaging and Bioengineering, the Canadian Institutes of Health Research, and private-sector contributions through the Foundation for the National Institutes of Health (FNIH), including generous contributions from the following: AbbVie; Alzheimer’s Association; Alzheimer’s Drug Discovery Foundation; Araclon Biotech; BioClinica, Inc.; Biogen; Bristol-Myers Squibb Company; CereSpir, Inc.; Cogstate; Eisai Inc.; Elan Pharmaceuticals, Inc.; Eli Lilly and Company; EuroImmun; F. Hoffmann-La Roche Ltd and its affiliated company Genentech, Inc.; Fujirebio; GE Healthcare; IXICO Ltd.; Janssen Alzheimer Immunotherapy Research & Development, LLC.; Johnson & Johnson Pharmaceutical Research & Development LLC.; Lumosity; Lundbeck; Merck & Co., Inc.; Meso Scale Diagnostics, LLC.; NeuroRx Research; Neurotrack Technologies; Novartis Pharmaceuticals Corporation; Pfizer Inc.; Piramal Imaging; Servier; Takeda Pharmaceutical Company; and Transition Therapeutics. We also gratefully acknowledge the HABS-HD study partners and their families, whose participation made this work possible. We are grateful to the Duke/UNC ADRC and AD-DECODE participants and their families.

