## SupplementaryMaterial for "Calibration-Aware and Interpretable Graph Learning for Multi-Cohort Diffusion Connectome Brain-Age Modeling"

**Supplementary Table Legends**

**Supplementary Table S1. MRI acquisition parameters by cohort.** Cohort-specific structural and diffusion MRI acquisition characteristics are summarized for ADNI, ADRC, HABS-HD, and AD-DECODE. ADNI data were acquired across multiple sites and scanner platforms and therefore had greater acquisition heterogeneity than the other cohorts, including both single-shell and multishell diffusion protocols. ADRC, HABS-HD, and AD-DECODE used comparatively standardized cohort-specific diffusion protocols. Across 347 ADRC diffusion datasets examined, all contained two b=0 volumes and 25 diffusion-weighted directions at b=800 s/mm². Among 491 HABS-HD acquisitions, 490 contained 12 b=0 volumes and 192 diffusion-weighted volumes at b=1000 s/mm². Among 53 non-backup AD-DECODE acquisitions with available gradient information, 52 contained three b=0 volumes and 21 diffusion-weighted directions at b=1000 s/mm². Acquisition heterogeneity was retained as an intrinsic component of cross-cohort model evaluation rather than harmonized at the raw-image/connectome level. DWI, diffusion-weighted imaging; FOV, field of view; MPRAGE, magnetization-prepared rapid gradient echo; MUSE, multiplexed sensitivity encoding; TE, echo time; TI, inversion time; TR, repetition time.

**Supplementary Table S2. Feature-set definitions and graph-aligned sample availability. A,** Predefined input configurations used for cohort-specific brain-age modeling. **B,** Graph-aligned sample availability for raw out-of-fold (OOF) model evaluation and full-cohort prediction. OOF graph sessions and participants denote the cognitively normal training and cross-validation sample used for each cohort and feature-set configuration. Prediction-ready full-cohort graph sessions denote graph-aligned observations available for full-cohort prediction and downstream validation, subject to analysis-specific feature and outcome availability. Cross-validation was grouped by participant to prevent leakage across repeated sessions. HABS-HD included 487 graph-aligned diffusion-weighted imaging/connectome sessions in the cohort summary; 486 passed the final prediction-readiness quality-control criteria used for full-cohort model outputs. Model-performance analyses used the OOF samples reported here. APOE, apolipoprotein E; BMI, body mass index; BP, blood pressure; DWI, diffusion-weighted imaging; FA, fractional anisotropy.

**Supplementary Table S3. Pooled raw out-of-fold brain-age prediction performance across cohorts.** Raw OOF predicted brain age was pooled across ADNI, ADRC, HABS-HD, and AD-DECODE for each feature-set model. Performance is summarized using mean absolute error (MAE), root mean squared error (RMSE), coefficient of determination ($R^2$), and Pearson correlation between predicted and chronological age, with participant-cluster bootstrap 95% confidence intervals. Models are ranked by pooled MAE. Full − CV denotes the full multimodal feature set excluding cardiovascular variables. Because pooled metrics are influenced by cohort sample size, composition, and age distribution, within-cohort performance was used as the primary evaluation framework.

**Supplementary Table S4. Across-cohort feature-set rank summary.** Feature-set models were ranked across ADNI, ADRC, HABS-HD, and AD-DECODE using the average of their within-cohort ranks from the raw OOF prediction analysis. Within-cohort ranks were based on lower MAE, lower RMSE, higher $R^2$, and higher Pearson correlation between predicted and chronological age. Ties in mean within-cohort rank were resolved using lower mean MAE, lower mean RMSE, higher mean $R^2$, and higher mean Pearson $r$, in that order. Full − CV denotes the full multimodal feature set excluding cardiovascular variables. This table summarizes relative feature-set robustness across cohorts and complements the primary within-cohort performance analysis.

**Supplementary Table S5. Hallmark gene-set enrichment analysis of PC17 in AD-DECODE.** Preranked gene-set enrichment analysis used the MSigDB Hallmark 2020 library. ES, enrichment score; NES, normalized enrichment score; FDR, false-discovery rate; FWER, family-wise error rate. Positive NES values indicate enrichment among positive PC17 gene loadings, corresponding to the lower-cBAG direction because imaging-only cBAG was negatively associated with PC17. Negative NES values indicate enrichment among negative PC17 loadings and therefore the higher-cBAG direction. At FDR $q < 0.05$, the lower-cBAG direction was enriched for heme-metabolism and pancreas-beta-cell signatures, whereas the higher-cBAG direction was enriched for interferon-alpha response. These analyses support biological interpretation of the latent PC17 transcriptomic component.

**Supplementary Table S6A. Imaging-only graph neural network and classical baseline performance by cohort.** Raw OOF performance is shown for the imaging-only graph neural network (GNN) and three conventional machine-learning baselines using a vectorized representation comprising connectome edge attributes, global graph metrics, and summary statistics of node-level imaging features. Models were evaluated within cohort. Lower MAE and RMSE and higher $R^2$ and Pearson correlation indicate better performance. GNN rows are shown in bold**. Supplementary Table S6B. Participant-clustered paired permutation comparisons between the GNN and classical baselines.** Paired comparisons used overlapping OOF predictions and 10,000 participant-clustered permutations. ΔMAE is baseline MAE minus GNN MAE; positive values favor the GNN. Two-sided permutation *P* values are reported for differences in absolute error and squared error. These analyses indicate broadly similar predictive performance across model classes rather than uniform superiority of the GNN.

**Supplementary Table S7. Medial temporal lobe and Alzheimer's disease–relevant exact-edge SHAP contributors.** The 28 prespecified medial temporal lobe (MTL) and Alzheimer's disease (AD)-relevant structural connectome edges included in the targeted enrichment analysis are listed in descending order of cross-cohort scaled SHAP importance. Global rank denotes the position among all 1,816 ranked exact-edge contributors, with smaller values indicating greater importance. Top percentile represents the corresponding position within the complete ranked edge distribution. This table provides the exact anatomical edge set used in the enrichment analyses summarized in Supplementary Table S8.

**Supplementary Table S8. Enrichment of MTL/AD-relevant exact-edge SHAP contributors.** The prespecified MTL/AD edge set was compared with the background distribution of all ranked exact-edge contributors using a one-sided Mann–Whitney U test and empirical permutation testing with 10,000 random edge sets of equal size. The complete selected-edge list is provided in Supplementary Table S7, allowing statistical enrichment results to be linked directly to individual anatomical connections.

**Supplementary Table S9. External versus within-cohort transferability summary.** Imaging-only cross-cohort transferability was summarized by comparing diagonal within-cohort out-of-fold (OOF) reference cells with off-diagonal external train–test combinations across ADNI, ADRC, HABS-HD, and AD-DECODE. Raw predicted-age metrics quantify chronological-age prediction performance, whereas corrected brain-age gap (cBAG) metrics characterize residual calibration after the prespecified age-bias correction. For MAE, RMSE, and mean absolute cBAG, lower values indicate better performance. For $R^2$ and Pearson r, higher values indicate better performance. For cBAG–age slope, values closer to zero indicate better calibration. Best and worst cells were identified according to these metric-specific criteria. No best or worst cell was assigned to the within-cohort cBAG–age slope because all values were effectively zero after correction. This table provides the quantitative summary corresponding to the complete transfer grid shown in Supplementary Table S10.

**Supplementary Table S10. Imaging-only transfer-grid metrics before and after cBAG correction.** Complete imaging-only train–test transfer metrics are shown for all 16 cohort combinations. Diagonal cells represent within-cohort OOF reference performance, whereas off-diagonal cells represent external transfer without retraining or recalibration. Raw prediction metrics quantify chronological-age prediction accuracy. Mean cBAG, mean absolute cBAG, and the residual cBAG-age slope characterize calibration after the prespecified age-bias correction. Age-informed recalibrated prediction metrics are intentionally omitted because they require target-cohort age information and therefore do not represent independent external-validation performance.

**Supplementary Table S11. Cell-wise post-hoc recalibration sensitivity metrics for cross-cohort transferability.** Cell-wise imaging-only transfer performance is shown before recalibration, after intercept-only recalibration, and after linear recalibration for all 16 training-cohort × testing-cohort combinations. Diagonal cells represent within-cohort OOF reference estimates, whereas off-diagonal cells represent external transfer without model retraining. Across the 12 external-transfer cells, median MAE decreased from 17.39 years before recalibration to 9.75 years after intercept-only recalibration and 8.22 years after linear recalibration. Median $R^2$ increased from −6.78 to −0.13 and 0.03, respectively, whereas median Pearson correlation remained unchanged at 0.171. Because affine recalibration alters prediction offset and scale but not participant ordering, Pearson correlation is identical across recalibration strategies. Recalibration parameters were estimated and evaluated within the same target cohort using chronological age and therefore represent a diagnostic sensitivity analysis rather than independent external-validation performance.

**Supplementary Figure Legends**

**Supplementary Figure S1. Cross-validation, age distribution, and training quality control across cohorts. A,** Chronological-age distributions for cognitively normal participants included in model training and participant-grouped cross-validation in ADNI, ADRC, HABS-HD, and AD-DECODE. **B,** Fold-level MAE calculated from raw OOF predicted age before age-bias correction across five predefined feature-set configurations in each cohort. Points represent individual held-out folds; boxes indicate the median and interquartile range, with whiskers extending to 1.5 times the interquartile range. **C,** Chronological-age distributions across the five participant-grouped cross-validation folds for the imaging-only configuration. Embedded boxplots indicate the median and interquartile range. **D,** Imaging-only GNN training loss across epochs. Curves show the cohort-specific mean training mean squared error after five-epoch centered smoothing, and shaded bands indicate 95% confidence intervals across folds. Training loss was computed on age targets standardized within each training fold and is expressed on a standardized-age scale rather than in years squared. **E,** Fold-level raw OOF validation MAE for the imaging-only model. **F,** Fold-level slopes from regressions of uncorrected brain-age gap on chronological age before age-bias correction across cohorts and feature-set configurations. Negative slopes indicate regression-to-the-mean age bias in raw brain-age gap.

**Supplementary Figure S2. Out-of-fold raw predicted age versus chronological age for the imaging-only model.** Scatter plots show chronological age versus raw OOF predicted brain age for the imaging-only GNN in each cohort. Each point represents one participant/session-level OOF prediction. Dashed blue lines show fitted linear associations between chronological and predicted age, and dotted orange lines show the identity line corresponding to perfect prediction. Insets report sample size, Pearson correlation, $R^2$, MAE, and RMSE. Raw predicted age was associated with chronological age in all cohorts, although fitted slopes were shallower than the identity line, consistent with regression-to-the-mean and supporting subsequent brain-age-gap correction before downstream validation.

**Supplementary Figure S3. Age-dependence diagnostics for raw and corrected brain-age gap across feature-set models.** Scatter plots show the association between chronological age and brain-age-gap measures pooled across feature-set models within each cohort. Columns show raw brain-age gap, fold-wise corrected brain-age gap, and OOF-global corrected brain-age gap; rows show cohorts. Each point represents a model-level participant/session prediction. Dashed lines show linear fits, and horizontal dotted lines indicate zero brain-age gap. Raw brain-age gap showed strong negative age dependence, whereas correction reduced this association. OOF-global corrected brain-age gap showed near-zero association with chronological age across cohorts, supporting its use as the primary age-corrected brain-age phenotype for downstream biological and longitudinal validation.

**Supplementary Figure S4. Extended longitudinal ΔcBAG associations across available ADNI, HABS-HD, and pooled endpoints.** Heatmaps show the number of complete longitudinal pairs and Pearson correlations for ΔcBAG associations across vascular, cognitive, imaging, and network endpoints. The forest plot summarizes all available longitudinal Δ associations with 95% confidence intervals, and endpoint Δ distributions are shown after z-scoring. These analyses were interpreted descriptively because endpoint availability, sample size, and effect direction varied across cohorts.

**Supplementary Figure S5. Longitudinal stability of brain-age metrics. A,** Annualized absolute longitudinal change in cBAG, cBAG centile, corrected predicted age, and predicted-age centile. Annualized change was calculated as absolute baseline-to-follow-up change divided by follow-up interval. Median annualized absolute ΔcBAG was 0.36 years/year in ADNI and 0.97 years/year in HABS-HD; median annualized absolute ΔcBAG-centile change was 3.31 and 5.21 centile points/year, respectively. **B,** Longitudinal preservation of corrected predicted age and predicted-age centiles. This analysis is shown as a reference because predicted age is expected to be more stable than cBAG owing to its strong dependence on chronological age.

**Supplementary Figure S6. Raw versus within-cohort scaled SHAP rankings for the imaging-only model.** Raw mean absolute SHAP rankings and within-cohort scaled SHAP rankings were compared across four SHAP feature classes in the imaging-only model. SHAP values were summarized across ADNI, ADRC, HABS-HD, and AD-DECODE for contributors present in all four cohorts. **A–B,** Global graph-feature rankings using raw mean absolute SHAP values (**A**) and within-cohort scaled SHAP values (**B**). **C–D,** Node-level feature rankings using raw (**C**) and scaled (**D**) SHAP values. **E–F,** Edge-derived regional rankings using raw (**E**) and scaled (**F**) SHAP values, where regional values summarize incident edge-level attributions. **G–H,** Exact-edge rankings using raw (**G**) and scaled (**H**) SHAP values; exact edges were restricted to features present in all four cohorts and supported by at least 100 graph sessions across cohorts. Scaled SHAP was calculated by dividing each feature's mean absolute SHAP value by the cohort-specific sum of mean absolute SHAP values within the same feature class before averaging across cohorts. Global graph-feature rankings were stable across raw and scaled analyses, whereas node-level, edge-derived regional, and exact-edge rankings showed partial reordering after scaling. This sensitivity analysis supports the use of within-cohort scaled SHAP values for the primary cross-cohort analysis while retaining raw SHAP values as an absolute-magnitude reference.

**Supplementary Figure S7. Extended anatomical visualization of SHAP contributors from the imaging-only brain-age model. A,** Top 30 node-level SHAP contributors, showing the anatomical distribution of the highest-ranking regional imaging features. **B,** Top 30 edge-derived regional contributors, in which exact-edge SHAP values were summarized at the regional level to identify brain regions participating most consistently in highly influential connectome edges. **C,** Top 30 exact-edge SHAP contributors, illustrating the highest-ranking structural connectome edges contributing to brain-age prediction. **D,** Targeted MTL/AD-relevant exact-edge map, highlighting connections involving the hippocampus, entorhinal cortex, parahippocampal cortex, amygdala, and related regions. Together, these complementary visualizations provide progressively finer anatomical interpretation of model attribution, from regional importance to individual connectome edges, while showing that medial temporal circuitry is represented among the highest-ranking exact-edge SHAP features.

**Supplementary Figure S8. Cross-cohort transferability before recalibration.** Cross-cohort transferability was evaluated by training the imaging-only brain-age model in one cohort and testing it in another across ADNI, ADRC, HABS-HD, and AD-DECODE. Rows indicate training cohorts and columns indicate testing cohorts. Diagonal cells represent within-cohort OOF reference performance, whereas off-diagonal cells represent external transfer without retraining or recalibration. Heatmaps summarize raw prediction error (MAE and RMSE), prediction correlation (Pearson *r*), explained variance ($R^2$), mean absolute cBAG, and residual cBAG-age slope for every train–test combination. External-transfer performance varied substantially across cohorts, with several off-diagonal combinations showing increased prediction error, reduced explained variance, and residual calibration bias relative to within-cohort reference models. These analyses illustrate the magnitude of cohort shift before recalibration and motivate the calibration-aware framework developed in the main manuscript.

**Supplementary Figure S9. Effect of post-hoc recalibration on cross-cohort transferability.** Cross-cohort transferability was re-evaluated after intercept-only and linear recalibration using target-cohort chronological age. Rows indicate training cohorts and columns indicate testing cohorts. Diagonal cells represent within-cohort OOF reference performance, whereas off-diagonal cells represent external transfer. Panels compare raw, intercept-recalibrated, and linear-recalibrated performance using MAE and $R^2$. Post-hoc recalibration substantially reduced prediction error and improved explained variance across most external-transfer combinations, demonstrating that external performance reflects both systematic calibration mismatch and preservation of subject-level predictive information. Although recalibration markedly improved MAE and $R^2$, Pearson correlation remained unchanged, indicating that calibration and discrimination represent complementary but distinct dimensions of cross-cohort transportability. Because recalibration uses target-cohort chronological age, these analyses characterize calibration sensitivity rather than independent external-validation performance.

**Supplementary Table S1. MRI acquisition parameters by cohort.**

| Cohort | Scanner/platform | Structural T1-weighted acquisition | Diffusion-weighted acquisition | Diffusion sampling |
|---|---|---|---|---|
| **ADNI** | 3 T; Siemens, GE, and Philips; multiple scanner models and sites | ADNI3 structural MRI protocols; acquisition parameters varied across scanner manufacturer and site | Conventional and multiband axial DTI; nearly all acquisitions used 2-mm slice thickness; TR, TE, matrix, phase encoding, and readout parameters varied across implementations | Predominantly single-shell (b=1000) s/mm² with approximately 30–48 diffusion-weighted directions and 1–7 (b=0) volumes; subset used multishell (b=500/1000/2000) s/mm² |
| **ADRC** | 3 T GE SIGNA UHP | 3D BRAVO/MPRAGE; TR/TE/TI = 2132.8/3.2/900 ms; flip angle = 8°; 256 × 256 matrix | Uniform single-shell DWI; paired reverse-phase-encoded images available for susceptibility-distortion correction | 27 volumes: 2 (b=0) + 25 directions at (b=800) s/mm² |
| **HABS-HD** | 3 T Siemens; MAGNETOM Vida and Skyra | 3D sagittal MPRAGE; TR/TE/TI = 2300/2.98/900 ms; flip angle = 9°; 1-mm slice thickness; 240 × 240 reconstruction matrix | AXIAL DTI mddw_64_SMS4; TE = 92 ms; nominal TR = 4000 ms; flip angle = 90°; 2.5-mm slice thickness in 489/491 acquisitions | Predominantly 204 volumes: 12 (b=0) + 192 diffusion-weighted volumes at (b=1000) s/mm² |
| **AD-DECODE** | 3 T GE SIGNA UHP | 3D axial MPRAGE; TR/TE/TI = 2193.6/3.036/900 ms; flip angle = 8°; 256 × 256 matrix; 1-mm isotropic resolution | 2D four-shot MUSE; TR/TE = 14,624/60.6 ms; flip angle = 90°; 256 × 256 matrix; 256 × 256-mm FOV; 1-mm reconstructed resolution | Predominantly 24 volumes: 3 (b=0) + 21 directions at (b=1000) s/mm² |

Cohort-specific structural and diffusion MRI acquisition characteristics are summarized for ADNI, ADRC, HABS-HD, and AD-DECODE. ADNI data were acquired across multiple sites and scanner platforms and therefore had greater acquisition heterogeneity than the other cohorts, including both single-shell and multishell diffusion protocols. ADRC, HABS-HD, and AD-DECODE used comparatively standardized cohort-specific diffusion protocols. Across 347 ADRC diffusion datasets examined, all contained two b=0 volumes and 25 diffusion-weighted directions at b=800 s/mm². Among 491 HABS-HD acquisitions, 490 contained 12 b=0 volumes and 192 diffusion-weighted volumes at b=1000 s/mm². Among 53 non-backup AD-DECODE acquisitions with available gradient information, 52 contained three b=0 volumes and 21 diffusion-weighted directions at b=1000 s/mm². Acquisition heterogeneity was retained as an intrinsic component of cross-cohort model evaluation rather than harmonized at the raw-image/connectome level. DWI, diffusion-weighted imaging; FOV, field of view; MPRAGE, magnetization-prepared rapid gradient echo; MUSE, multiplexed sensitivity encoding; TE, echo time; TI, inversion time; TR, repetition time.

**Supplementary Table S2. Feature-set definitions and graph-aligned sample availability.**

**Supplementary Table S2A. Feature-set definitions.**

| Feature set | Table label | Required input blocks |
|---|---|---|
| Imaging only | Imaging only | DWI/connectome graph inputs; regional FA and volume measures where available; graph-theoretic metrics |
| Imaging + demographic/genetic covariates | Imaging + demo. | Imaging-only inputs plus sex and APOE genotype |
| Imaging + biomarkers | Imaging + biomarkers | Imaging-only inputs plus available cohort-specific biomarkers or transcriptomic principal components |
| Full model without cardiovascular variables | Full − CV | Imaging-only inputs plus sex, APOE genotype, BMI, and available biomarker/transcriptomic features; excludes systolic blood pressure, diastolic blood pressure, and pulse |
| Full model | Full | Imaging-only inputs plus sex, APOE genotype, BMI, cardiovascular variables, and available biomarker/transcriptomic features |

APOE, apolipoprotein E; BMI, body mass index; BP, blood pressure; DWI, diffusion-weighted imaging; FA, fractional anisotropy.

**Supplementary Table S2B. Graph-aligned out-of-fold sample availability by cohort and feature set.**

| Cohort | Feature set | OOF graph sessions, n | OOF participants, n | Prediction-ready full-cohort graph sessions, n | Notes |
|---|---|---|---|---|---|
| **ADNI** | | | | | |
| | Imaging only | 233 | 140 | 316 | Cognitively normal OOF training/CV sample |
| | Imaging + demo. | 233 | 140 | 316 | Same OOF sample retained |
| | Imaging + biomarkers | 233 | 140 | 316 | Same OOF sample retained |
| | Full − CV | 233 | 140 | 316 | Cardiovascular variables excluded |
| | Full | 233 | 140 | 316 | Full multimodal feature set |
| **ADRC** | | | | | |
| | Imaging only | 145 | 145 | 204 | Cognitively normal OOF training/CV sample |
| | Imaging + demo. | 145 | 145 | 204 | Same OOF sample retained |
| | Imaging + biomarkers | 145 | 145 | 204 | Same OOF sample retained |
| | Full − CV | 145 | 145 | 204 | Cardiovascular variables excluded |
| | Full | 145 | 145 | 204 | Full multimodal feature set |
| **HABS-HD** | | | | | |
| | Imaging only | 394 | 272 | 486 | Cognitively normal OOF training/CV sample |
| | Imaging + demo. | 394 | 272 | 486 | Same OOF sample retained |
| | Imaging + biomarkers | 394 | 272 | 486 | Same OOF sample retained |
| | Full − CV | 394 | 272 | 486 | Cardiovascular variables excluded |
| | Full | 394 | 272 | 486 | Full multimodal feature set |
| **AD-DECODE** | | | | | |
| | Imaging only | 79 | 79 | 86 | Cognitively normal OOF training/CV sample |
| | Imaging + demo. | 79 | 79 | 86 | Same OOF sample retained |
| | Imaging + biomarkers | 79 | 79 | 86 | Same OOF sample retained |
| | Full − CV | 79 | 79 | 86 | Cardiovascular variables excluded |
| | Full | 79 | 79 | 86 | Full multimodal feature set |

**A,** Predefined input configurations used for cohort-specific brain-age modeling. **B,** Graph-aligned sample availability for raw out-of-fold (OOF) model evaluation and full-cohort prediction. OOF graph sessions and participants denote the cognitively normal training and cross-validation sample used for each cohort and feature-set configuration. Prediction-ready full-cohort graph sessions denote graph-aligned observations available for full-cohort prediction and downstream validation, subject to analysis-specific feature and outcome availability. Cross-validation was grouped by participant to prevent leakage across repeated sessions. HABS-HD included 487 graph-aligned diffusion-weighted imaging/connectome sessions in the cohort summary; 486 passed the final prediction-readiness quality-control criteria used for full-cohort model outputs. Model-performance analyses used the OOF samples reported here. APOE, apolipoprotein E; BMI, body mass index; BP, blood pressure; DWI, diffusion-weighted imaging; FA, fractional anisotropy.

**Supplementary Table S3. Pooled raw out-of-fold brain-age prediction performance across cohorts.**

| Rank | Model | N sessions | N participants | MAE [95% CI] | RMSE [95% CI] | $R^2$ [95% CI] | Pearson r [95% CI] |
|---|---|---|---|---|---|---|---|
| 1 | Full − CV | 851 | 636 | 5.676 [5.286, 6.036] | 7.589 [7.062, 8.068] | 0.623 [0.570, 0.669] | 0.790 [0.757, 0.819] |
| 2 | Imaging + biomarkers | 851 | 636 | 5.693 [5.306, 6.084] | 7.747 [7.165, 8.395] | 0.607 [0.539, 0.660] | 0.780 [0.739, 0.813] |
| 3 | Full | 851 | 636 | 5.727 [5.362, 6.093] | 7.639 [7.163, 8.109] | 0.618 [0.569, 0.660] | 0.787 [0.756, 0.816] |
| 4 | Imaging + demo. | 851 | 636 | 5.897 [5.525, 6.301] | 7.898 [7.421, 8.431] | 0.592 [0.533, 0.638] | 0.771 [0.733, 0.802] |
| 5 | Imaging only | 851 | 636 | 5.937 [5.553, 6.331] | 7.974 [7.413, 8.526] | 0.584 [0.514, 0.642] | 0.764 [0.722, 0.803] |

Raw out-of-fold predicted brain age was pooled across ADNI, ADRC, HABS-HD, and AD-DECODE for each feature-set model. Performance is summarized using mean absolute error (MAE), root mean squared error (RMSE), coefficient of determination ($R^2$), and Pearson correlation between predicted and chronological age, with participant-cluster bootstrap 95% confidence intervals. Models are ranked by pooled MAE. Full − CV denotes the full multimodal feature set excluding cardiovascular variables. The pooled analysis metrics are influenced by cohort sample size, cohort composition, and differences in age distribution; within-cohort performance was therefore used as the primary evaluation framework.

**Supplementary Table S4. Across-cohort feature-set rank summary.**

| Across-cohort rank | Model | N cohorts | Mean within-cohort rank | Mean MAE | Mean RMSE | Mean $R^2$ | Mean Pearson r |
|---|---|---|---|---|---|---|---|
| 1 | Full − CV | 4 | 2.50 | 6.679 | 8.565 | 0.285 | 0.538 |
| 2 | Full | 4 | 2.50 | 6.886 | 8.661 | 0.276 | 0.534 |
| 3 | Imaging + biomarkers | 4 | 3.00 | 6.643 | 8.755 | 0.256 | 0.493 |
| 4 | Imaging only | 4 | 3.00 | 6.897 | 8.925 | 0.231 | 0.476 |
| 5 | Imaging + demo. | 4 | 4.00 | 6.909 | 8.859 | 0.240 | 0.487 |

Feature-set models were ranked across ADNI, ADRC, HABS-HD, and AD-DECODE using the average of their within-cohort ranks from the raw out-of-fold prediction analysis. Within-cohort ranks were based on lower mean absolute error (MAE), lower root mean squared error (RMSE), higher coefficient of determination ($R^2$), and higher Pearson correlation between predicted and chronological age. Ties in mean within-cohort rank were resolved using lower mean MAE, lower mean RMSE, higher mean $R^2$, and higher mean Pearson r. Full − CV denotes the full multimodal feature set excluding cardiovascular variables. This table summarizes relative feature-set robustness across cohorts and supplements the primary within-cohort performance analysis.

**Supplementary Table S5. Hallmark gene-set enrichment analysis of PC17 in AD-DECODE.**

| Hallmark term | ES | NES | Nominal p | FDR q | FWER p | Tag fraction | Gene fraction | Leading-edge genes |
|---|---|---|---|---|---|---|---|---|
| Heme metabolism | 0.525603 | 2.558871 | 0.001 | 0.001 | 0.001 | 79/179 | 15.34% | MXI1; AQP3; TRIM58; TAL1; SNCA; ANK1; TRIM10; SLC25A38; CTNS; CPOX; KLF1; EPB42; ALAS2; GLRX5; RBM38; SLC4A1; HBB; SELENBP1; SLC6A8; DMTN; MYL4; BCAM; TENT5C; ADIPOR1; BACH1; BPGM; ICAM4; TSPAN5; CTSE; TNS1; SPTB; BNIP3L; BTG2; RANBP10; HBQ1; MKRN1; E2F2; PPP2R5B; MAP2K3; GMPS; BLVRB; SLC25A37; GYPC; HDGF; OSBP2; HAGH; SPTA1; FOXO3; PDZK1IP1; GATA1; MPP1; FBXO9; AHSP; OPTN; EPOR; FBXO7; TOP1; RNF123; CTSB; PIGQ; H1-0; BSG; RAD23A; EPB41; CLCN3; CLIC2; XK; UROD; CAST; RIOK3; MARCHF8; SLC66A2; CA1; ARHGEF12; GCLC; SMOX; TMCC2; MFHAS1; BMP2K |
| Pancreas beta cells | 0.617438 | 1.839383 | 0.00464 | 0.008435 | 0.012 | 10/16 | 16.64% | NEUROD1; VDR; ELP4; FOXO1; PAK3; MAFB; SRPRB; DPP4; SEC11A; SRP9 |
| Interferon alpha response | −0.422940 | −1.758320 | 0.001 | 0.014473 | 0.019 | 53/91 | 34.49% | MOV10; RTP4; OASL; PARP12; STAT2; TRIM21; SAMD9; IRF1; IFI44L; IFIT3; SLC25A28; HERC6; CMTR1; IRF9; UBE2L6; ISG15; DHX58; RSAD2; IFIT2; BST2; IRF2; MX1; SAMD9L; PSME2; PARP14; CNP; LAP3; ADAR; TMEM140; BATF2; OAS1; GBP2; SELL; IFI35; HELZ2; OGFR; GBP4; PSME1; TAP1; B2M; ISG20; UBA7; DDX60; PSMB9; IRF7; PARP9; SP110; IFITM1; USP18; WARS1; TRAFD1; CCRL2; TRIM25 |
| TNF-alpha signaling via NF-kB | 0.322186 | 1.549138 | 0.001 | 0.069356 | 0.136 | 43/156 | 18.64% | PTX3; IL23A; TUBB2A; CD44; TSC22D1; SPHK1; MYC; NR4A1; TRAF1; IRS2; BTG2; EIF1; MAP2K3; TNIP1; FUT4; KLF6; SERPINE1; ATP2B1; IL7R; DUSP5; BIRC2; MARCKS; CD69; EGR1; SAT1; SERPINB8; DENND5A; DRAM1; PNRC1; SOCS3; JUNB; KLF2; CDKN1A; RHOB; TNIP2; GCH1; IFIH1; YRDC; ZC3H12A; KLF10; CEBPB; IFNGR2; FOS |
| Interferon gamma response | −0.337280 | −1.545310 | 0.003082 | 0.086460 | 0.207 | 81/184 | 30.97% | CD40; SLAMF7; RTP4; CASP7; OASL; IL18BP; PARP12; CMKLR1; STAT2; TRIM21; IRF1; SRI; IFI44L; IFIT3; APOL6; IRF5; GBP6; SLC25A28; HERC6; SERPING1; STAT1; ZBP1; CMTR1; IL2RB; RBCK1; GZMA; RNF213; PSMA2; IRF9; UBE2L6; ISG15; DHX58; RSAD2; IFIT2; BST2; IRF2; MX1; GPR18; SAMD9L; CCL5; PSME2; IL10RA; IFIT1; PARP14; RIPK1; MT2A; LAP3; ADAR; BATF2; C1R; LYSMD2; MX2; TNFAIP3; KLRK1; HLA-A; IFI35; HELZ2; OGFR; CD274; GBP4; TNFAIP2; PSME1; CIITA; TAP1; TOR1B; IL15RA; SSPN; OAS2; XAF1; NOD1; LATS2; CD38; B2M; ISG20; PSMB10; DDX60; PSMB9; IRF7; SPPL2A; LCP2; SP110 |
| KRAS signaling down | 0.320061 | 1.353635 | 0.039506 | 0.203006 | 0.497 | 20/75 | 15.04% | PRODH; KCND1; TENT5C; NOS1; RYR1; CCDC106; BTG2; SLC5A5; KRT1; GP2; TFCP2L1; ZBTB16; MAST3; LYPD3; ENTPD7; GP1BA; SERPINA10; KCNMB1; CDKAL1; GAMT |
| G2-M checkpoint | 0.274268 | 1.326340 | 0.021978 | 0.214394 | 0.577 | 41/159 | 17.64% | ESPL1; SLC7A5; CUL5; GINS2; MYC; HMGB3; TFDP1; CDC20; GSPT1; MYBL2; CENPF; E2F2; SMC4; E2F1; CUL4A; PTTG1; HMGA1; MARCKS; KIF5B; TOP1; CTCF; E2F3; TPX2; CCNT1; PRPF4B; PRMT5; BIRC5; BUB3; CBX1; XPO1; HIF1A; NEK2; NUP98; SQLE; MNAT1; INCENP; ATRX; TRA2B; H2AX; TACC3; HUS1 |
| KRAS signaling up | 0.292634 | 1.357693 | 0.028329 | 0.248135 | 0.489 | 24/130 | 10.91% | EMP1; FLT4; ZNF639; PCSK1N; SPON1; AMMECR1; PLEK2; TRAF1; C3AR1; BPGM; RETN; NAP1L2; WNT7A; PLVAP; ANKH; RELN; EPB41L3; GADD45G; GYPC; IL7R; HKDC1; EPHB2; MAFB; SCN1B |
| IL-2/STAT5 signaling | 0.263799 | 1.268398 | 0.053221 | 0.288603 | 0.754 | 35/161 | 15.44% | EMP1; TNFRSF18; CD44; CCR4; SPRED2; TGM2; MYC; TRAF1; MUC1; BMPR2; BCL2L1; RNH1; RRAGD; KLF6; GATA1; NFKBIZ; PUS1; PIM1; LTB; CDKN1C; DENND5A; ENO3; SMPDL3A; MYO1C; GPX4; AHCY; SH3BGRL2; FLT3LG; NT5E; IRF4; CASP3; RHOB; LCLAT1; ICOS; PTRH2 |

Preranked gene-set enrichment analysis used the MSigDB Hallmark 2020 library. ES, enrichment score; NES, normalized enrichment score; FDR, false-discovery rate; FWER, family-wise error rate. Positive NES values indicate enrichment among positive PC17 gene loadings, which correspond to the lower-cBAG direction because imaging-only cBAG was negatively associated with PC17. Negative NES values indicate enrichment among negative PC17 loadings and therefore the higher-cBAG direction. At FDR $q < 0.05$, the lower-cBAG direction was enriched for heme metabolism and pancreas beta-cell signatures, whereas the higher-cBAG direction was enriched for interferon alpha response. The transcriptomic analysis aids biological interpretation of the latent PC17 component.

### Supplementary Table S6A. Imaging-only graph neural network and classical baseline performance by cohort.

| Cohort | Family | Model | N sessions | N participants | MAE | RMSE | R² | Pearson r |
|---|---|---|---|---|---|---|---|---|
| ADNI | Classical | Elastic net | 233 | 140 | 4.715 | 5.939 | 0.143 | 0.388 |
| **ADNI** | **GNN** | **GNN** | **233** | **140** | **4.724** | **5.883** | **0.159** | **0.399** |
| ADNI | Classical | Ridge | 233 | 140 | 5.016 | 6.548 | -0.042 | 0.316 |
| ADNI | Classical | XGBoost | 233 | 140 | 5.046 | 6.192 | 0.068 | 0.294 |
| ADRC | Classical | Ridge | 145 | 145 | 7.448 | 10.005 | 0.220 | 0.506 |
| ADRC | Classical | Elastic net | 145 | 145 | 7.509 | 10.036 | 0.215 | 0.503 |
| ADRC | Classical | XGBoost | 145 | 145 | 7.525 | 9.481 | 0.300 | 0.549 |
| **ADRC** | **GNN** | **GNN** | **145** | **145** | **7.967** | **10.678** | **0.112** | **0.360** |
| HABS | Classical | XGBoost | 394 | 272 | 5.031 | 6.364 | 0.385 | 0.624 |
| HABS | Classical | Elastic net | 394 | 272 | 5.111 | 6.414 | 0.375 | 0.615 |
| **HABS** | **GNN** | **GNN** | **394** | **272** | **5.143** | **6.579** | **0.342** | **0.586** |
| HABS | Classical | Ridge | 394 | 272 | 5.635 | 7.825 | 0.070 | 0.488 |
| **AD DECODE** | **GNN** | **GNN** | **79** | **79** | **9.753** | **12.561** | **0.310** | **0.558** |
| AD DECODE | Classical | XGBoost | 79 | 79 | 9.920 | 12.421 | 0.326 | 0.571 |
| AD DECODE | Classical | Ridge | 79 | 79 | 10.264 | 12.963 | 0.266 | 0.524 |
| AD DECODE | Classical | Elastic net | 79 | 79 | 10.725 | 13.464 | 0.208 | 0.483 |

Raw OOF performance is shown for the imaging-only graph neural network (GNN) and three conventional machine-learning baselines using a vectorized representation of the connectome edge attributes and global graph metrics. Models were evaluated within cohort. Lower MAE and RMSE and higher $R^2$ and Pearson correlation indicate better performance. GNN rows are shown in bold.

### Supplementary Table S6B. Participant-clustered paired permutation comparisons between the GNN and classical baselines.

| Cohort | Baseline model | N sessions | N participants | Baseline MAE | GNN MAE | ΔMAE | Paired P (MAE) | Baseline RMSE | GNN RMSE | Paired P (MSE) |
|---|---|---|---|---|---|---|---|---|---|---|
| ADNI | Elastic net | 233 | 140 | 4.715 | 4.724 | -0.008 | 0.5170 | 5.939 | 5.883 | 0.3878 |
| ADNI | Ridge | 233 | 140 | 5.016 | 4.724 | 0.292 | 0.1522 | 6.548 | 5.883 | 0.0292 |
| ADNI | XGBoost | 233 | 140 | 5.046 | 4.724 | 0.322 | 0.0416 | 6.192 | 5.883 | 0.1261 |
| ADRC | Ridge | 145 | 145 | 7.448 | 7.967 | -0.519 | 0.4222 | 10.005 | 10.678 | 0.4454 |
| ADRC | Elastic net | 145 | 145 | 7.509 | 7.967 | -0.458 | 0.4711 | 10.036 | 10.678 | 0.4606 |
| ADRC | XGBoost | 145 | 145 | 7.525 | 7.967 | -0.442 | 0.3810 | 9.481 | 10.678 | 0.0740 |
| HABS | XGBoost | 394 | 272 | 5.031 | 5.143 | -0.112 | 0.4546 | 6.364 | 6.579 | 0.1939 |
| HABS | Elastic net | 394 | 272 | 5.111 | 5.143 | -0.032 | 0.8351 | 6.414 | 6.579 | 0.4736 |
| HABS | Ridge | 394 | 272 | 5.635 | 5.143 | 0.492 | 0.2520 | 7.825 | 6.579 | 0.1847 |
| AD DECODE | XGBoost | 79 | 79 | 9.920 | 9.753 | 0.167 | 0.8205 | 12.421 | 12.561 | 0.8477 |
| AD DECODE | Ridge | 79 | 79 | 10.264 | 9.753 | 0.510 | 0.5165 | 12.963 | 12.561 | 0.6858 |
| AD DECODE | Elastic net | 79 | 79 | 10.725 | 9.753 | 0.972 | 0.2462 | 13.464 | 12.561 | 0.4435 |

Paired comparisons used overlapping OOF predictions and 10,000 participant-clustered permutations. ΔMAE is baseline MAE minus GNN MAE; positive values favor the GNN. Two-sided permutation *P* values are reported for differences in absolute error and squared error. These analyses indicate broadly similar predictive performance across model classes rather than uniform superiority of the GNN.

### Supplementary Table S7. Medial temporal lobe and Alzheimer's disease–relevant exact-edge SHAP contributors.

| Target-set rank | Exact edge | Mean scaled \|SHAP\| | Global rank | Top percentile (%) |
|---|---|---|---|---|
| 1 | ctx-lh-fusiform -- Left-Hippocampus | 0.000238 | 7 | 0.39 |
| 2 | ctx-lh-parahippocampal -- Left-Hippocampus | 0.000237 | 8 | 0.44 |
| 3 | ctx-rh-lingual -- Right-Hippocampus | 0.000236 | 9 | 0.50 |
| 4 | ctx-rh-parahippocampal -- Right-Hippocampus | 0.000236 | 10 | 0.55 |
| 5 | ctx-rh-fusiform -- Right-Hippocampus | 0.000235 | 12 | 0.66 |
| 6 | Left-Amygdala -- Left-Hippocampus | 0.000232 | 16 | 0.88 |
| 7 | Right-Amygdala -- Right-Hippocampus | 0.000228 | 22 | 1.21 |
| 8 | ctx-lh-entorhinal -- Left-Hippocampus | 0.000227 | 24 | 1.32 |
| 9 | ctx-rh-entorhinal -- Right-Hippocampus | 0.000225 | 32 | 1.76 |
| 10 | ctx-lh-lingual -- Left-Hippocampus | 0.000224 | 36 | 1.98 |
| 11 | ctx-rh-precuneus -- Right-Hippocampus | 0.000217 | 65 | 3.58 |
| 12 | ctx-lh-entorhinal -- ctx-lh-medialorbitofrontal | 0.000216 | 70 | 3.85 |
| 13 | Left-Thalamus-Proper -- Left-Hippocampus | 0.000201 | 164 | 9.03 |
| 14 | Right-Thalamus-Proper -- Right-Hippocampus | 0.000201 | 172 | 9.47 |
| 15 | Left-Putamen -- Left-Hippocampus | 0.000178 | 420 | 23.13 |
| 16 | Right-Hippocampus -- Right-Putamen | 0.000176 | 459 | 25.28 |
| 17 | ctx-lh-superiortemporal -- Right-Hippocampus | 0.000176 | 468 | 25.77 |
| 18 | Right-Hippocampus -- ctx-lh-middletemporal | 0.000171 | 564 | 31.06 |
| 19 | ctx-rh-superiorfrontal -- Left-Hippocampus | 0.000169 | 589 | 32.43 |
| 20 | ctx-rh-entorhinal -- ctx-lh-medialorbitofrontal | 0.000159 | 815 | 44.88 |
| 21 | ctx-rh-fusiform -- Left-Hippocampus | 0.000159 | 816 | 44.93 |
| 22 | ctx-lh-insula -- Right-Hippocampus | 0.000150 | 1007 | 55.45 |
| 23 | ctx-rh-insula -- Left-Hippocampus | 0.000148 | 1062 | 58.48 |
| 24 | Right-Hippocampus -- ctx-lh-supramarginal | 0.000147 | 1088 | 59.91 |
| 25 | ctx-lh-parstriangularis -- Left-Hippocampus | 0.000145 | 1128 | 62.11 |
| 26 | ctx-rh-entorhinal -- Right-Caudate | 0.000140 | 1247 | 68.67 |
| 27 | ctx-rh-entorhinal -- Right-Thalamus-Proper | 0.000137 | 1311 | 72.19 |
| 28 | ctx-rh-entorhinal -- ctx-rh-postcentral | 0.000133 | 1365 | 75.17 |

The 28 prespecified medial temporal lobe (MTL) and Alzheimer's disease (AD)-relevant structural connectome edges included in the targeted enrichment analysis are listed in descending order of cross-cohort scaled SHAP importance. Global rank denotes the position among all 1,816 ranked exact-edge contributors, with smaller values indicating greater importance. Top percentile represents the corresponding percentile within the complete ranked edge distribution. Top percentile is calculated from the global rank, with smaller percentages indicating placement nearer the top of the distribution. This table provides the exact edge set used for the enrichment analyses summarized in Supplementary Table S8.

**Supplementary Table S8. Enrichment of MTL/AD-relevant exact-edge SHAP contributors.**

| Measure | Result |
|---|---|
| Total globally ranked exact edges | 1816 |
| Prespecified MTL/AD edges matched | 28 |
| Prespecified MTL/AD edges missing | 0 |
| Mean importance, MTL/AD edges | 1.91e-04 |
| Mean importance, background edges | 1.53e-04 |
| Median importance, MTL/AD edges | 1.90e-04 |
| Median importance, background edges | 1.55e-04 |
| Mann-Whitney U | 37484 |
| One-sided Mann-Whitney P | <0.0001 |
| Random edge sets | 10000 |
| Empirical P, mean importance | <0.0001 |
| Empirical P, median importance | <0.0001 |
| Median global rank | 296 |
| Best-worst global rank | 7-1365 |
| Selected edges in top 5% | 12/28 (42.9%) |

The prespecified MTL/AD edge set was compared with the background distribution of all ranked exact-edge contributors using a one-sided Mann-Whitney U test and empirical permutation testing with 10,000 random edge sets of equal size. The complete selected-edge list is provided in Supplementary Table S7, allowing statistical enrichment results to be linked directly to individual anatomical connections.

**Supplementary Table S9. External versus within-cohort transferability summary.**

| Comparison group | Metric | N cells | Minimum | Median | Mean | Maximum | Best cell | Worst cell |
|---|---|---|---|---|---|---|---|---|
| Within-cohort OOF | Raw MAE | 4 | 4.724 | 6.555 | 6.897 | 9.753 | ADNI→ADNI | AD-DECODE→AD-DECODE |
| Within-cohort OOF | Raw RMSE | 4 | 5.883 | 8.629 | 8.925 | 12.561 | ADNI→ADNI | AD-DECODE→AD-DECODE |
| Within-cohort OOF | Raw $R^2$ | 4 | 0.112 | 0.235 | 0.231 | 0.342 | HABS-HD→HABS-HD | ADRC→ADRC |
| Within-cohort OOF | Raw Pearson r | 4 | 0.360 | 0.479 | 0.476 | 0.586 | HABS-HD→HABS-HD | ADRC→ADRC |
| Within-cohort OOF | Mean \|cBAG\| | 4 | 1.793 | 3.288 | 3.366 | 5.097 | ADNI→ADNI | AD-DECODE→AD-DECODE |
| External transfer | Raw MAE | 12 | 8.821 | 17.394 | 30.462 | 80.573 | ADNI→HABS-HD | AD-DECODE→ADNI |
| External transfer | Raw RMSE | 12 | 10.799 | 22.677 | 38.373 | 81.294 | ADNI→HABS-HD | HABS-HD→ADRC |
| External transfer | Raw $R^2$ | 12 | -156.846 | -6.781 | -28.446 | -0.151 | ADRC→AD-DECODE | AD-DECODE→ADNI |
| External transfer | Raw Pearson r | 12 | -0.024 | 0.171 | 0.213 | 0.514 | ADRC→HABS-HD | HABS-HD→ADRC |
| External transfer | Mean \|cBAG\| | 12 | 2.333 | 8.176 | 19.819 | 65.559 | ADNI→HABS-HD | AD-DECODE→HABS-HD |
| External transfer | cBAG-age slope | 12 | -0.489 | -0.110 | -0.099 | 0.226 | ADRC→ADNI | HABS-HD→ADRC |

Imaging-only cross-cohort transferability was summarized by comparing diagonal within-cohort out-of-fold (OOF) reference cells with off-diagonal external train–test combinations across ADNI, ADRC, HABS-HD, and AD-DECODE. Raw predicted-age metrics quantify chronological-age prediction performance, whereas corrected brain-age gap (cBAG) metrics characterize residual calibration after the prespecified age-bias correction. For MAE, RMSE, and mean absolute cBAG, lower values indicate better performance. For $R^2$ and Pearson r, higher values indicate better performance. For cBAG–age slope, values closer to zero indicate better calibration. Best and worst cells were identified according to these metric-specific criteria. No best or worst cell was assigned to the within-cohort cBAG–age slope because all values were effectively zero after correction. This table provides the quantitative summary corresponding to the complete transfer grid shown in Supplementary Table S10.

**Supplementary Table S10. Imaging-only transfer-grid metrics before and after cBAG correction.**

| Training cohort | Testing cohort | Cell type | N | Raw MAE | Raw RMSE | Raw R² | Raw Pearson r | Mean cBAG | Mean \|cBAG\| | cBAG-age slope | Slope P |
|---|---|---|---|---|---|---|---|---|---|---|---|
| ADNI | ADNI | Within-cohort OOF | 233 | 4.724 | 5.883 | 0.159 | 0.399 | -0.000 | 1.793 | -0.000 | 1.0000 |
| ADNI | ADRC | External | 204 | 17.204 | 20.547 | -2.023 | 0.239 | 1.514 | 2.786 | -0.046 | 0.1259 |
| ADNI | HABS-HD | External | 486 | 8.821 | 10.799 | -0.609 | 0.319 | -0.260 | 2.333 | -0.034 | 0.0329 |
| ADNI | AD-DECODE | External | 86 | 29.954 | 33.780 | -3.444 | 0.224 | 9.668 | 9.668 | -0.119 | <0.0001 |
| ADRC | ADNI | External | 316 | 19.713 | 21.536 | -10.119 | 0.159 | -2.831 | 5.050 | -0.008 | 0.8928 |
| ADRC | ADRC | Within-cohort OOF | 145 | 7.967 | 10.678 | 0.112 | 0.360 | 0.000 | 3.711 | -0.000 | 1.0000 |
| ADRC | HABS-HD | External | 486 | 11.460 | 13.365 | -1.466 | 0.514 | -1.775 | 4.002 | 0.145 | <0.0001 |
| ADRC | AD-DECODE | External | 86 | 14.427 | 17.193 | -0.151 | 0.122 | 3.468 | 4.013 | -0.162 | <0.0001 |
| HABS-HD | ADNI | External | 316 | 16.426 | 23.817 | -12.599 | 0.179 | -7.106 | 12.029 | 0.226 | 0.1851 |
| HABS-HD | ADRC | External | 204 | 16.424 | 81.294 | < −20 | -0.024 | 3.773 | 14.323 | -0.489 | 0.3032 |
| HABS-HD | HABS-HD | Within-cohort OOF | 394 | 5.143 | 6.579 | 0.342 | 0.586 | 0.000 | 2.864 | 0.000 | 1.0000 |
| HABS-HD | AD-DECODE | External | 86 | 17.583 | 21.371 | -0.779 | 0.463 | 5.743 | 6.683 | -0.159 | <0.0001 |
| AD-DECODE | ADNI | External | 316 | 80.573 | 81.144 | < −20 | 0.163 | -61.993 | 61.993 | -0.100 | 0.1591 |
| AD-DECODE | ADRC | External | 204 | 55.125 | 56.718 | < −20 | 0.126 | -49.320 | 49.386 | -0.225 | <0.0001 |
| AD-DECODE | HABS-HD | External | 486 | 77.829 | 78.914 | < −20 | 0.074 | -65.559 | 65.559 | -0.217 | 0.0001 |
| AD-DECODE | AD-DECODE | Within-cohort OOF | 79 | 9.753 | 12.561 | 0.310 | 0.558 | -0.000 | 5.097 | 0.000 | 1.0000 |

Complete imaging-only train-test transfer metrics are shown for all 16 cohort combinations. Diagonal cells represent within-cohort out-of-fold reference performance, whereas off-diagonal cells represent external transfer without retraining or recalibration. Raw prediction metrics quantify chronological-age prediction accuracy. Mean corrected brain-age gap (cBAG), mean absolute cBAG, and the residual cBAG-age slope characterize calibration after the prespecified age-bias correction. Age-informed recalibrated prediction metrics are intentionally omitted because they require target-cohort age information and therefore do not represent independent external-validation performance.

**Supplementary Table S11. Cell-wise post-hoc recalibration sensitivity metrics for cross-cohort transferability.**

| Training cohort | Testing cohort | Cell type | N | MAE raw | MAE intercept | MAE linear | RMSE raw | RMSE intercept | RMSE linear | $R^2$ raw | $R^2$ intercept | $R^2$ linear | Pearson r |
|---|---|---|---|---|---|---|---|---|---|---|---|---|---|
| ADNI | ADNI | Within-cohort OOF | 233 | 4.724 | 4.725 | 4.715 | 5.883 | 5.883 | 5.882 | 0.159 | 0.159 | 0.159 | 0.399 |
| ADNI | ADRC | External transfer | 204 | 17.204 | 9.576 | 9.543 | 20.547 | 11.724 | 11.476 | -2.023 | 0.016 | 0.057 | 0.239 |
| ADNI | HABS-HD | External transfer | 486 | 8.821 | 6.553 | 6.539 | 10.799 | 8.077 | 8.066 | -0.609 | 0.099 | 0.102 | 0.319 |
| ADNI | AD-DECODE | External transfer | 86 | 29.954 | 13.320 | 13.234 | 33.780 | 15.668 | 15.617 | -3.444 | 0.044 | 0.050 | 0.224 |
| ADRC | ADNI | External transfer | 316 | 19.713 | 6.718 | 5.265 | 21.536 | 8.673 | 6.377 | -10.119 | -0.803 | 0.025 | 0.159 |
| ADRC | ADRC | Within-cohort OOF | 145 | 7.967 | 7.979 | 8.182 | 10.678 | 10.676 | 10.569 | 0.112 | 0.112 | 0.130 | 0.360 |
| ADRC | HABS-HD | External transfer | 486 | 11.460 | 5.846 | 5.786 | 13.365 | 7.366 | 7.302 | -1.466 | 0.251 | 0.264 | 0.514 |
| ADRC | AD-DECODE | External transfer | 86 | 14.427 | 13.322 | 13.333 | 17.193 | 15.905 | 15.904 | -0.151 | 0.015 | 0.015 | 0.122 |
| HABS-HD | ADNI | External transfer | 316 | 16.426 | 9.930 | 5.236 | 23.817 | 19.725 | 6.354 | -12.599 | -8.327 | 0.032 | 0.179 |
| HABS-HD | ADRC | External transfer | 204 | 16.424 | 18.900 | 9.852 | 81.294 | 80.675 | 11.813 | -46.326 | -45.609 | 0.001 | -0.024 |
| HABS-HD | HABS-HD | Within-cohort OOF | 394 | 5.143 | 5.139 | 5.137 | 6.579 | 6.579 | 6.574 | 0.342 | 0.342 | 0.343 | 0.586 |
| HABS-HD | AD-DECODE | External transfer | 86 | 17.583 | 12.008 | 11.794 | 21.371 | 14.303 | 14.205 | -0.779 | 0.203 | 0.214 | 0.463 |
| AD-DECODE | ADNI | External transfer | 316 | 80.573 | 7.529 | 5.214 | 81.144 | 9.617 | 6.372 | -156.846 | -1.217 | 0.027 | 0.163 |
| AD-DECODE | ADRC | External transfer | 204 | 55.125 | 10.659 | 9.774 | 56.718 | 13.347 | 11.722 | -22.037 | -0.276 | 0.016 | 0.126 |
| AD-DECODE | HABS-HD | External transfer | 486 | 77.829 | 9.370 | 6.887 | 78.914 | 13.040 | 8.488 | -84.952 | -1.347 | 0.006 | 0.074 |
| AD-DECODE | AD-DECODE | Within-cohort OOF | 79 | 9.753 | 9.785 | 9.777 | 12.561 | 12.552 | 12.552 | 0.310 | 0.311 | 0.312 | 0.558 |

Cell-wise imaging-only transfer performance is shown before recalibration, after intercept-only recalibration, and after linear recalibration for all 16 training-cohort × testing-cohort combinations. Diagonal cells represent within-cohort out-of-fold reference estimates, whereas off-diagonal cells represent external transfer without model retraining. Across the 12 external-transfer cells, median MAE decreased from 17.39 years before recalibration to 9.75 years after intercept-only recalibration and 8.22 years after linear recalibration. Median $R^2$ increased from −6.78 to −0.13 and 0.03, respectively, whereas median Pearson correlation remained unchanged at 0.171. Because affine recalibration alters prediction offset and scale but not participant ordering, Pearson correlation is identical across recalibration strategies. Recalibration parameters were estimated and evaluated within the same target cohort using chronological age and therefore represent a diagnostic sensitivity analysis rather than independent external-validation performance.

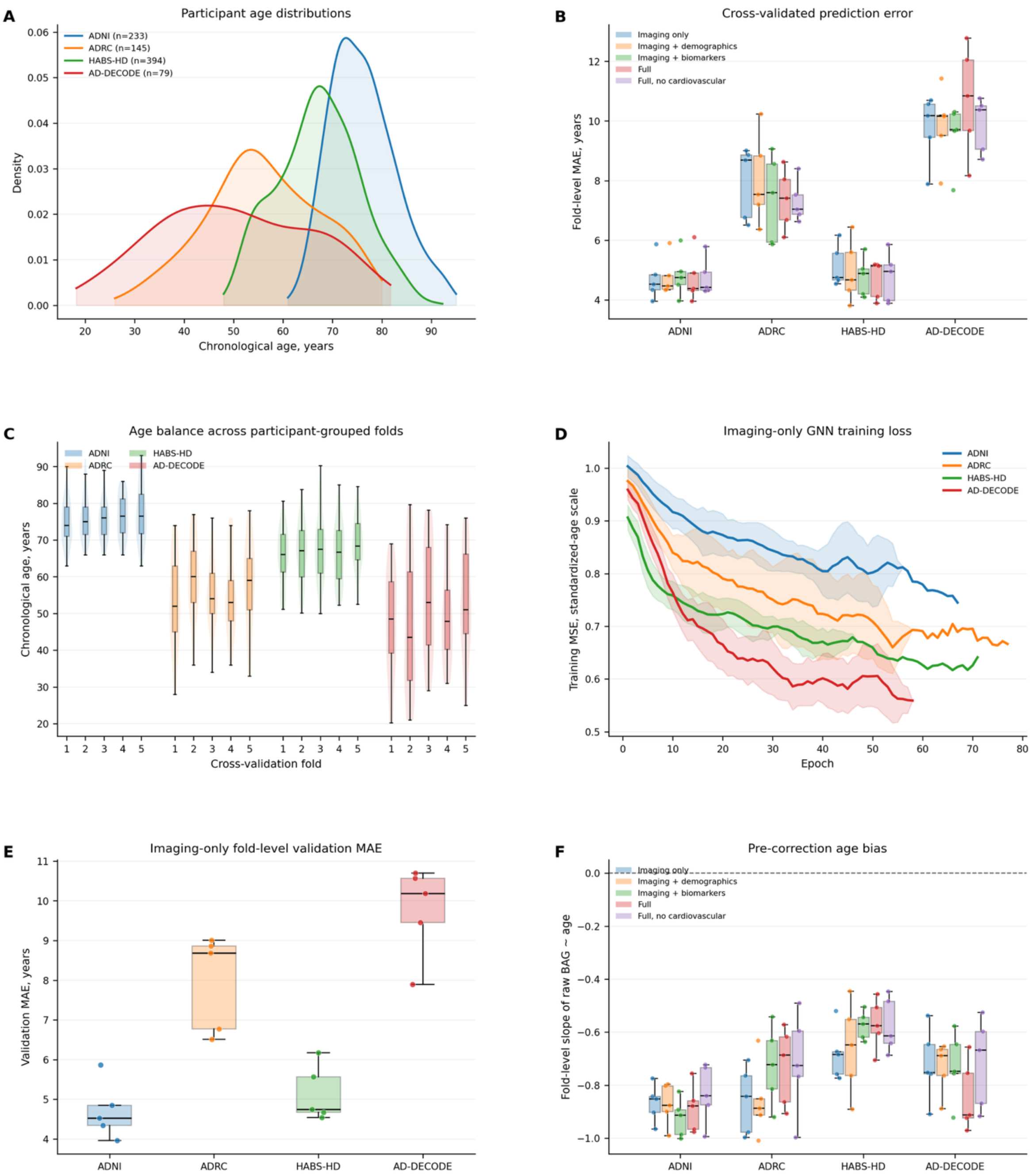


**Supplementary Figure S1. Cross-validation, age distribution, and training quality control across cohorts.** **A**, Chronological-age distributions for cognitively normal participants included in model training and participant-grouped cross-validation in ADNI, ADRC, HABS-HD, and AD-DECODE. **B**, Fold-level mean absolute error (MAE) calculated from raw out-of-fold predicted age before age-bias correction, across five predefined feature-set configurations in each cohort. Points represent individual held-out folds; boxes indicate the median and interquartile range, with whiskers extending to 1.5 times the interquartile range. **C**, Chronological-age distributions across the five participant-grouped cross-validation folds for the imaging-only configuration. The embedded boxplots indicate the median and interquartile range. **D**, Imaging-only graph neural network training loss across epochs. Curves show the cohort-specific mean training mean squared error after five-epoch centered smoothing, and shaded bands indicate 95% confidence intervals across folds. Training loss was computed on age targets standardized within each training fold and is expressed on a standardized-age scale rather than in years squared. **E**, Fold-level raw out-of-fold validation MAE for the imaging-only model. Points represent individual held-out folds; boxes indicate the median and interquartile range. **F**, Fold-level slopes from regressions of uncorrected brain-age gap on chronological age before age-bias correction, shown across cohorts and feature-set configurations. Negative slopes indicate regression-to-the-mean age bias in raw brain-age gap.

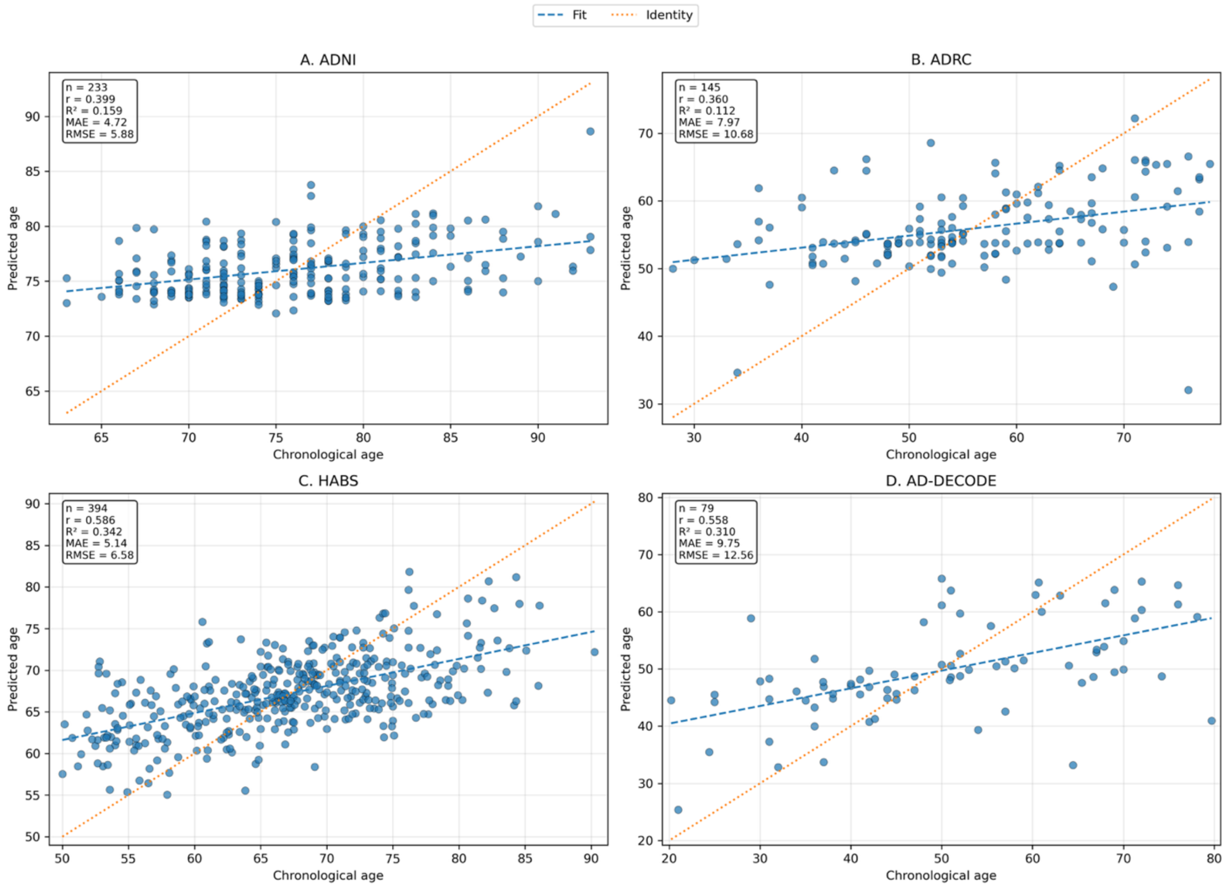


**Supplementary Figure S2. Out-of-fold raw predicted age versus chronological age for the imaging-only model.** Scatter plots show chronological age versus raw out-of-fold predicted brain age for the imaging-only graph neural network model in each cohort. Each point represents one participant/session-level out-of-fold prediction. Dashed blue lines show the fitted linear association between chronological and predicted age, and dotted orange lines show the identity line corresponding to perfect prediction. Insets report sample size, Pearson correlation coefficient, coefficient of determination ((R^2)), mean absolute error (MAE), and root mean squared error (RMSE). Raw predicted age was associated with chronological age in all cohorts, although fitted slopes were shallower than the identity line, consistent with regression-to-the-mean in raw brain-age prediction and supporting subsequent brain-age gap correction before downstream validation analyses.

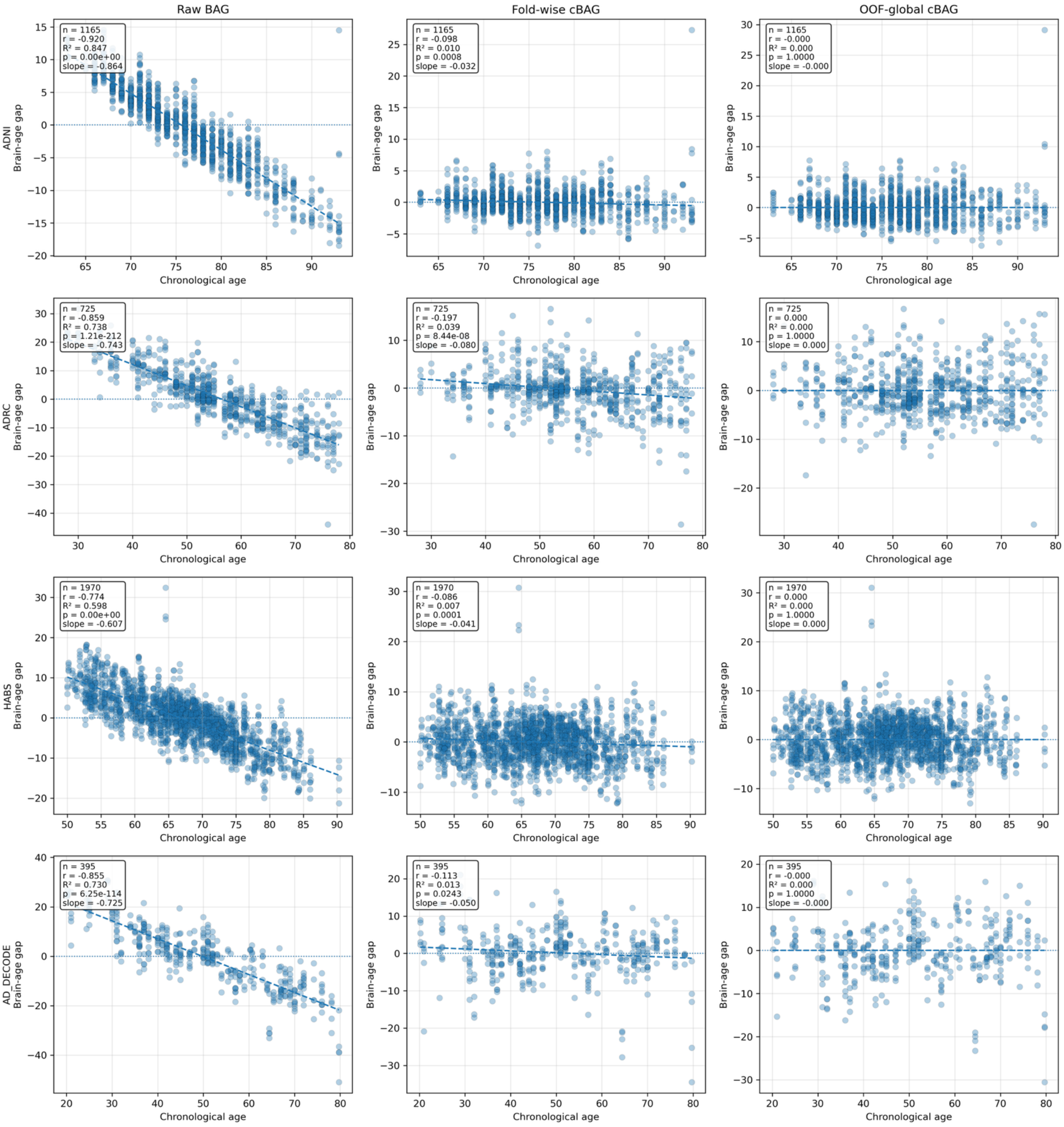


**Supplementary Figure S3. Age-dependence diagnostics for raw and corrected brain-age gap across feature-set models.** Scatter plots show the association between chronological age and brain-age gap measures pooled across feature-set models within each cohort. Columns show raw brain-age gap, fold-wise corrected brain-age gap, and OOF-global corrected brain-age gap. Rows show cohorts. Each point represents a model-level participant/session prediction. Dashed lines show linear fits, and horizontal dotted lines indicate zero brain-age gap. Raw brain-age gap showed strong negative age-dependence, whereas correction reduced this association. OOF-global corrected brain-age gap showed near-zero association with chronological age across cohorts, supporting its use as the primary age-corrected brain-age phenotype for downstream biological and longitudinal validation analyses.

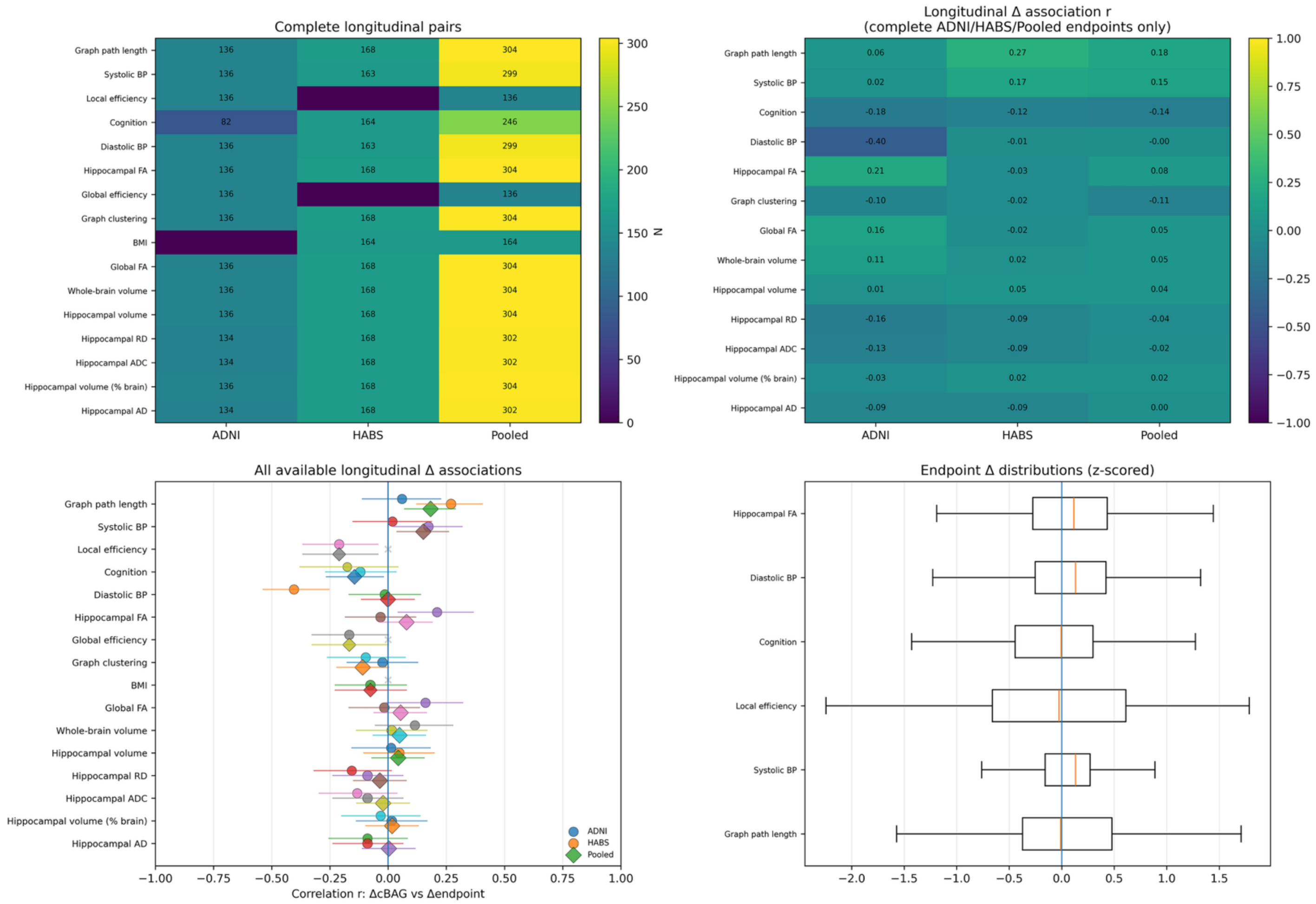


**Supplementary Figure S4. E**xtended longitudinal ΔcBAG associations across available ADNI, HABS-HD, and pooled endpoints. Heatmaps show the number of complete longitudinal pairs and Pearson correlation coefficients for ΔcBAG associations across vascular, cognitive, imaging, and network endpoints. The forest plot summarizes all available longitudinal Δ associations with 95% confidence intervals, and endpoint Δ distributions are shown after z-scoring. These analyses were interpreted descriptively because endpoint availability and effect directions varied across cohorts.

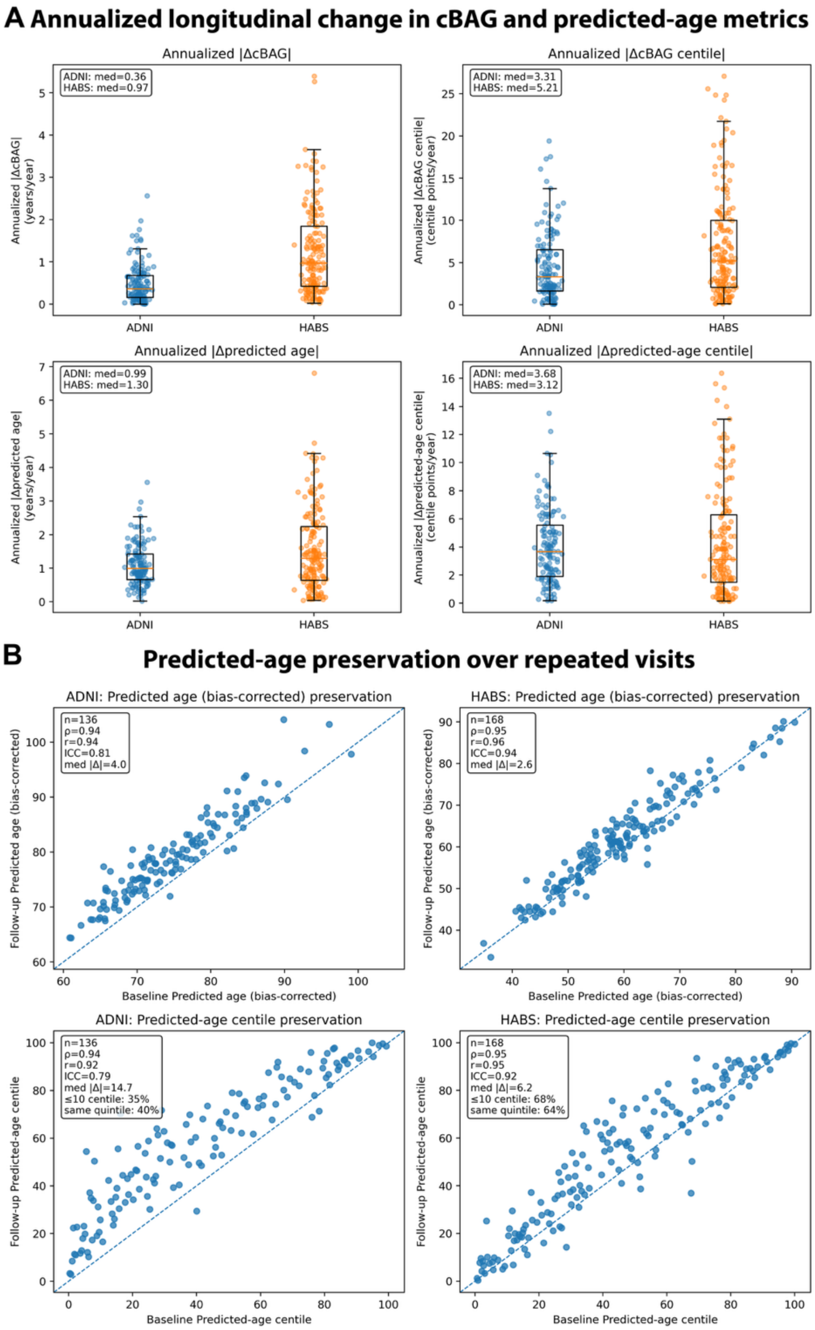


**Supplementary Figure S5. Longitudinal stability of brain-age metrics. A.** Annualized absolute longitudinal change in cBAG, cBAG centile, corrected predicted age, and predicted-age centile. Annualized change was calculated as absolute baseline-to-follow-up change divided by follow-up interval. Median annualized absolute ΔcBAG was 0.36 years/year in ADNI and 0.97 years/year in HABS-HD; median annualized absolute ΔcBAG-centile change was 3.31 and 5.21 centile points/year, respectively. **B,** longitudinal preservation of corrected predicted age and predicted-age centiles. This analysis is shown as a reference because predicted age is expected to be more stable than cBAG due to its strong dependence on chronological age.

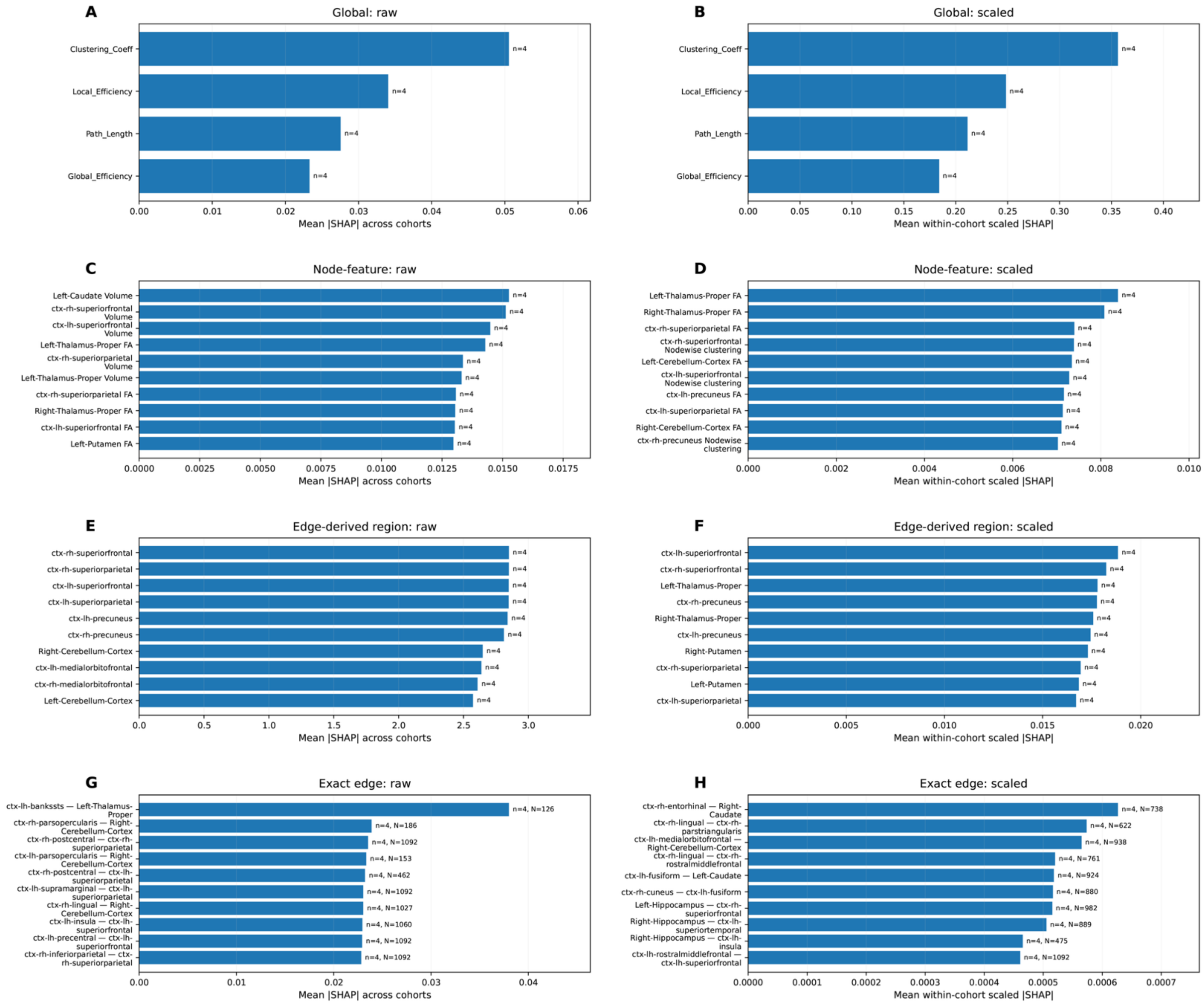


**Supplementary Figure S6. Raw versus within-cohort scaled SHAP rankings for the imaging-only model.** Raw mean absolute SHAP rankings and within-cohort scaled SHAP rankings were compared across four SHAP feature classes in the imaging-only model. SHAP values were summarized across ADNI, ADRC, HABS, and AD-DECODE for contributors present in all four cohorts. **A–B**, Global graph-feature rankings using raw mean absolute SHAP values (**A**) and within-cohort scaled SHAP values (**B**). **C–D**, Node-level feature rankings using raw (**C**) and within-cohort scaled (**D**) SHAP values. **E–F**, Edge-derived regional rankings using raw (**E**) and within-cohort scaled (**F**) SHAP values, where regional values summarize incident edge-level attributions. **G–H**, Exact-edge rankings using raw (**G**) and within-cohort scaled (**H**) SHAP values; exact edges were restricted to features present in all four cohorts and supported by at least 100 graph sessions across cohorts. Scaled SHAP was calculated by dividing each feature's mean absolute SHAP value by the cohort-specific sum of mean absolute SHAP values within the same SHAP feature class before averaging across cohorts. Global graph-feature rankings were stable across raw and scaled analyses, whereas node-level, edge-derived regional, and exact-edge rankings showed partial reordering after scaling. This sensitivity analysis supports the use of within-cohort scaled SHAP values for the primary cross-cohort analysis while retaining raw SHAP values as an absolute-magnitude reference.

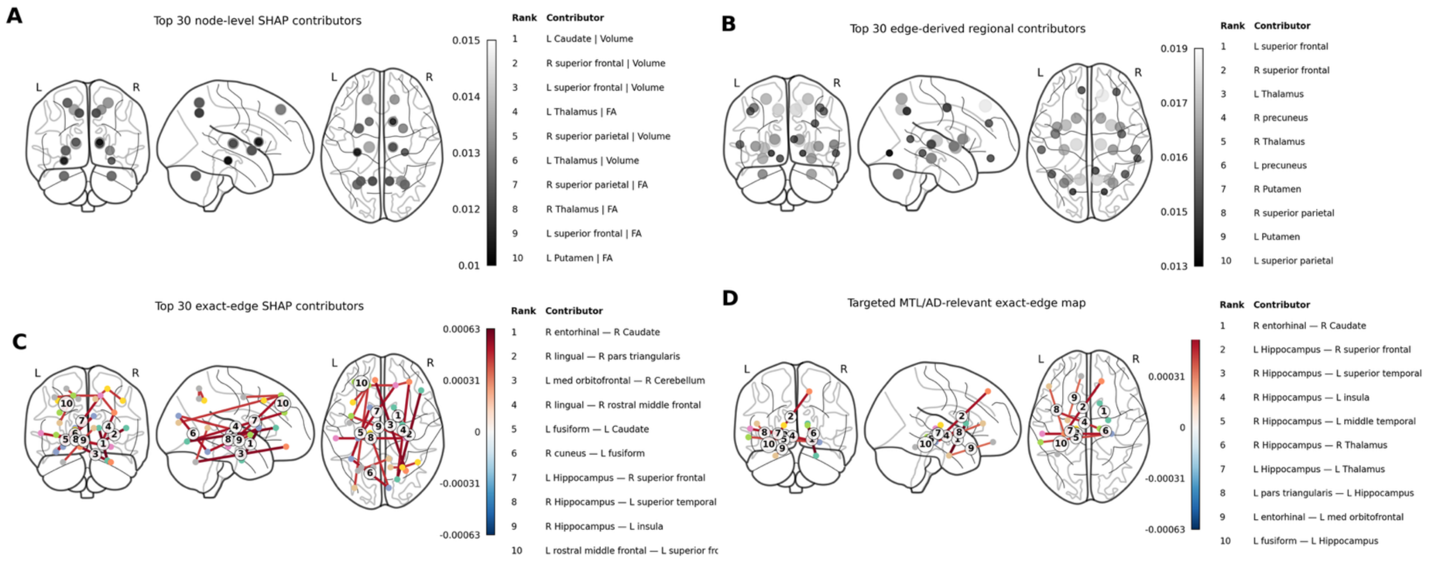


**Supplementary Figure S7. Extended anatomical visualization of SHAP contributors from the imaging-only brain-age model.** A, Top 30 node-level SHAP contributors, showing the anatomical distribution of the highest-ranking regional imaging features. B, Top 30 edge-derived regional contributors, in which exact-edge SHAP values were summarized at the regional level to identify brain regions participating most consistently in highly influential connectome edges. C, Top 30 exact-edge SHAP contributors, illustrating the highest-ranking structural connectome edges contributing to brain-age prediction. D, Medial temporal lobe subset of the highest-ranking exact-edge contributors, highlighting connections involving the hippocampus, entorhinal cortex, parahippocampal cortex, amygdala, and related regions to facilitate anatomical interpretation of the learned connectome. Together, these complementary visualizations provide progressively finer anatomical interpretation of model attribution, from regional importance to individual connectome edges, while showing that medial temporal circuitry is represented among the highest-ranking exact-edge SHAP features.

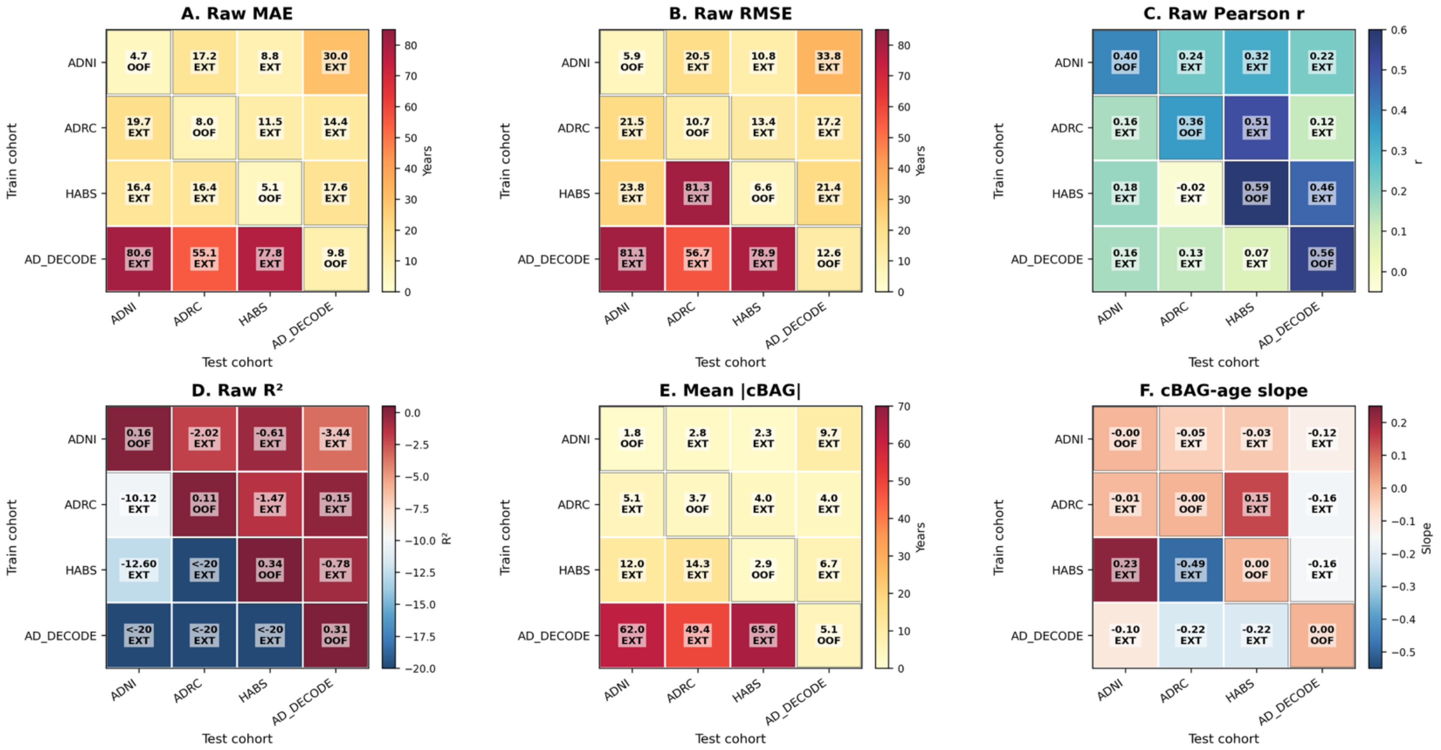


**Supplementary Figure S8. Cross-cohort transferability before recalibration.** Cross-cohort transferability was evaluated by training the imaging-only brain-age model in one cohort (rows) and testing it in another (columns) across ADNI, ADRC, HABS-HD, and AD-DECODE. Diagonal cells represent within-cohort out-of-fold reference performance, whereas off-diagonal cells represent external transfer without retraining or recalibration. Heatmaps summarize raw prediction error (MAE, RMSE), prediction correlation (Pearson *r*), explained variance ($R^2$), mean absolute corrected brain-age gap (|cBAG|), and residual cBAG-age slope for every train-test combination. External transfer performance varied substantially across cohorts, with several off-diagonal combinations exhibiting increased prediction error, reduced explained variance, and residual calibration bias relative to the corresponding within-cohort reference models. These analyses illustrate the magnitude of domain shift before recalibration and motivate the calibration-aware framework developed in the main manuscript.

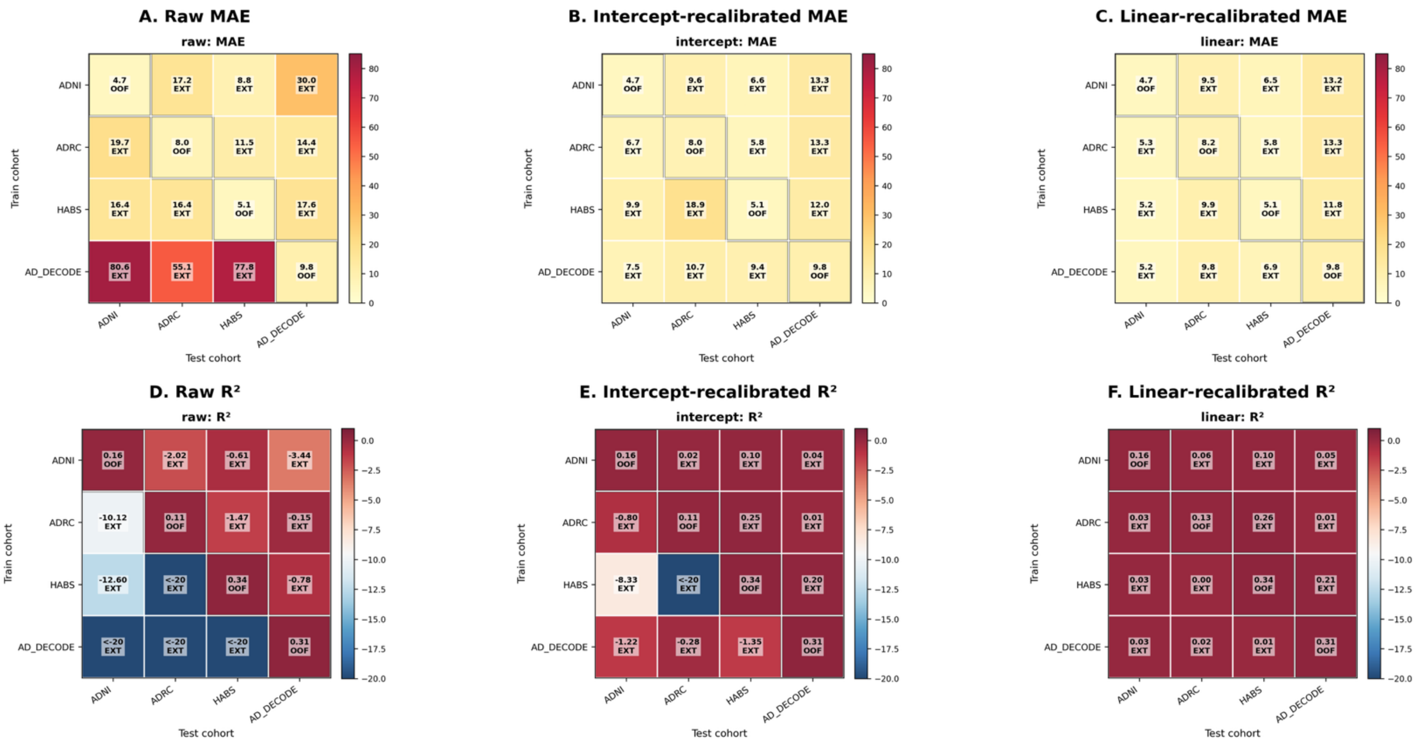

**Supplementary Figure S9. Effect of post-hoc recalibration on cross-cohort transferability.** Post-hoc recalibration substantially reduced prediction error and improved explained variance across most external-transfer combinations, demonstrating that external performance reflects both systematic calibration mismatch and preservation of subject-level predictive information. Although recalibration markedly improved MAE and $R^2$, Pearson correlation remained unchanged, indicating that calibration and discrimination represent complementary but distinct dimensions of cross-cohort transportability. Because recalibration uses target-cohort chronological age, these analyses characterize calibration sensitivity rather than independent external-validation performance.
